## Supplementary figures and images for "UNRAVELING CRP/cAMP-MEDIATED METABOLIC REGULATION IN *ESCHERICHIA COLI* PERSISTER CELLS"

### Figure 1 - figure supplement 1

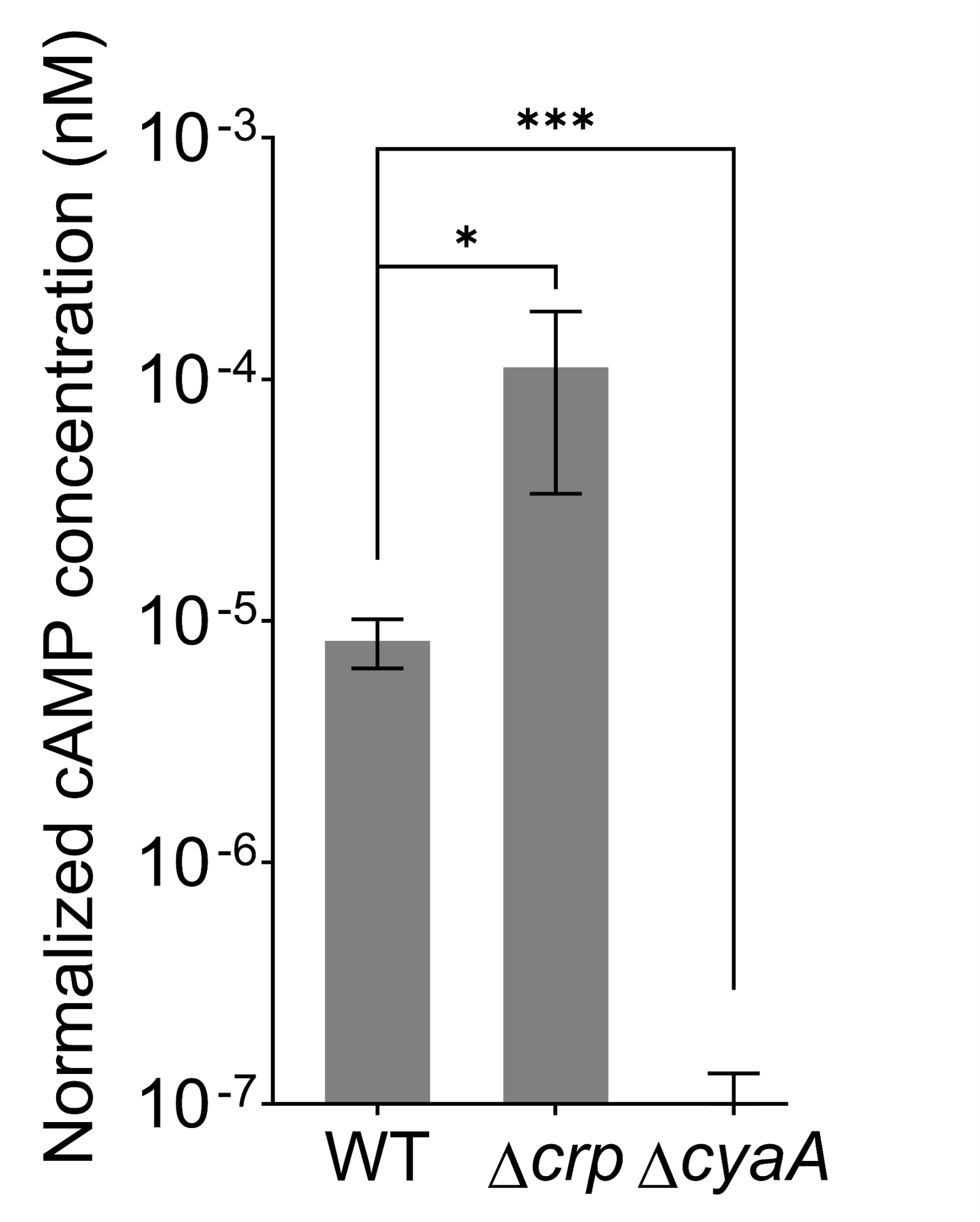

### Figure 1 - figure supplement 2

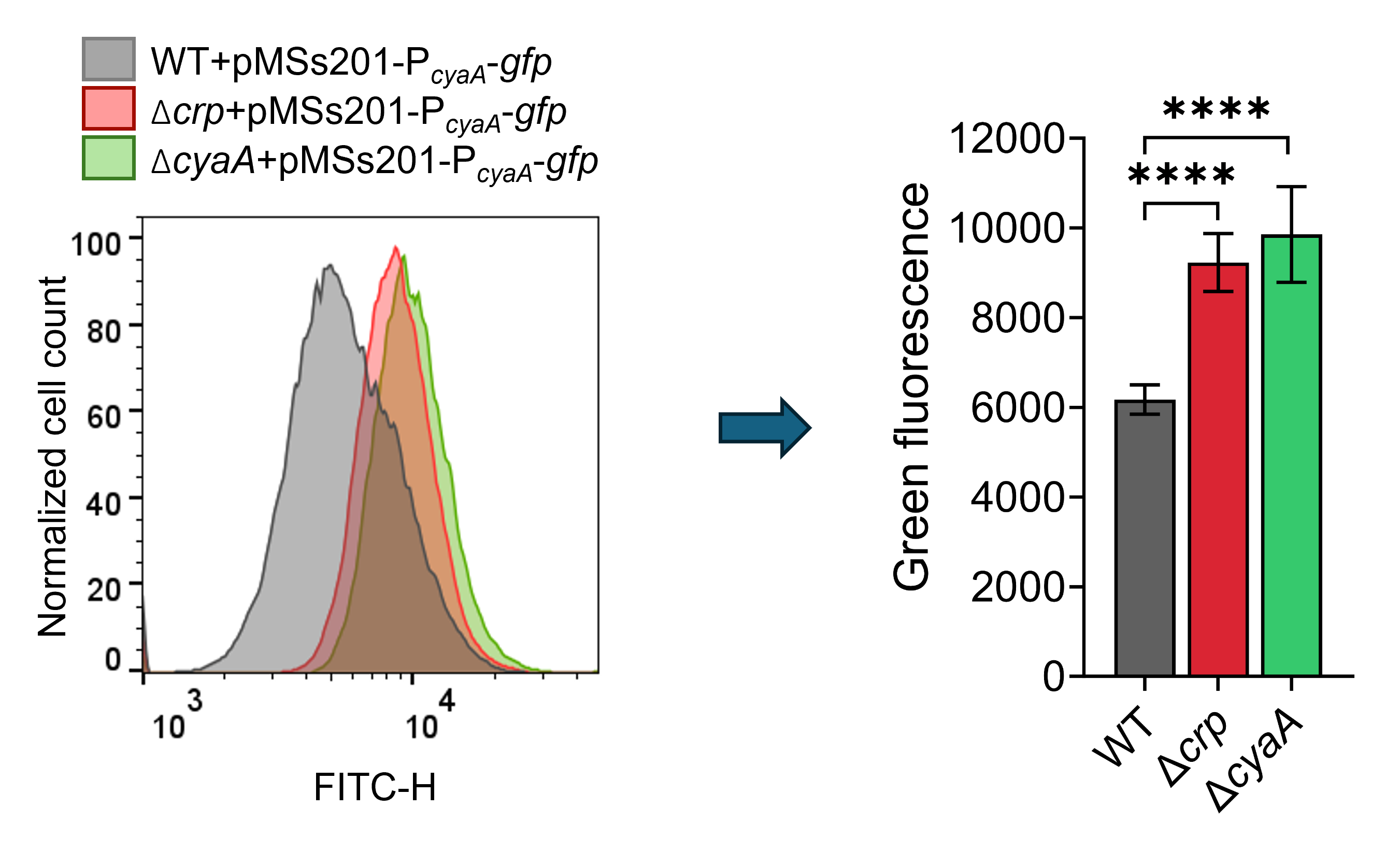

### Figure 1 - figure supplement 3

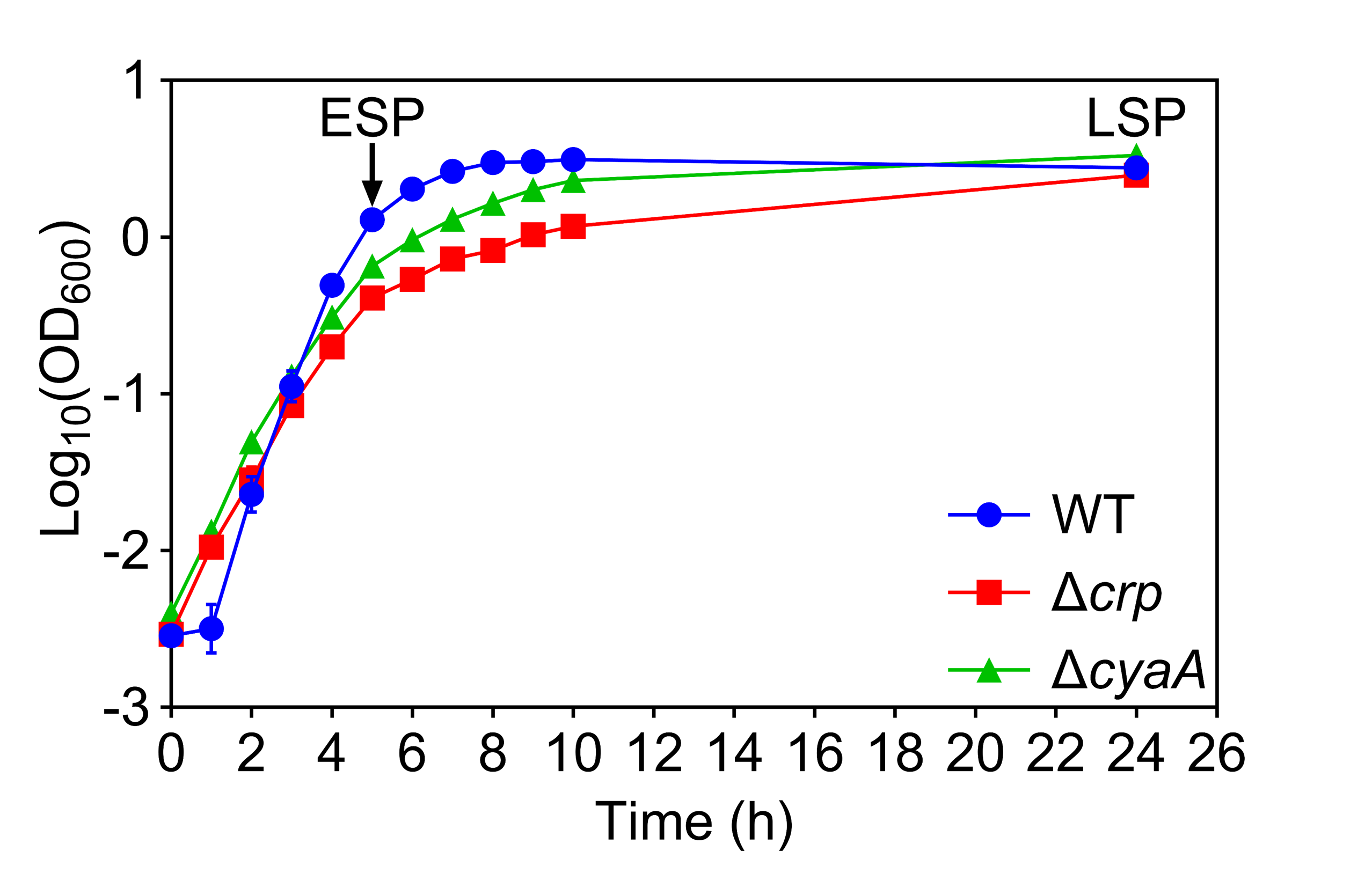

### Figure 1 - figure supplement 4

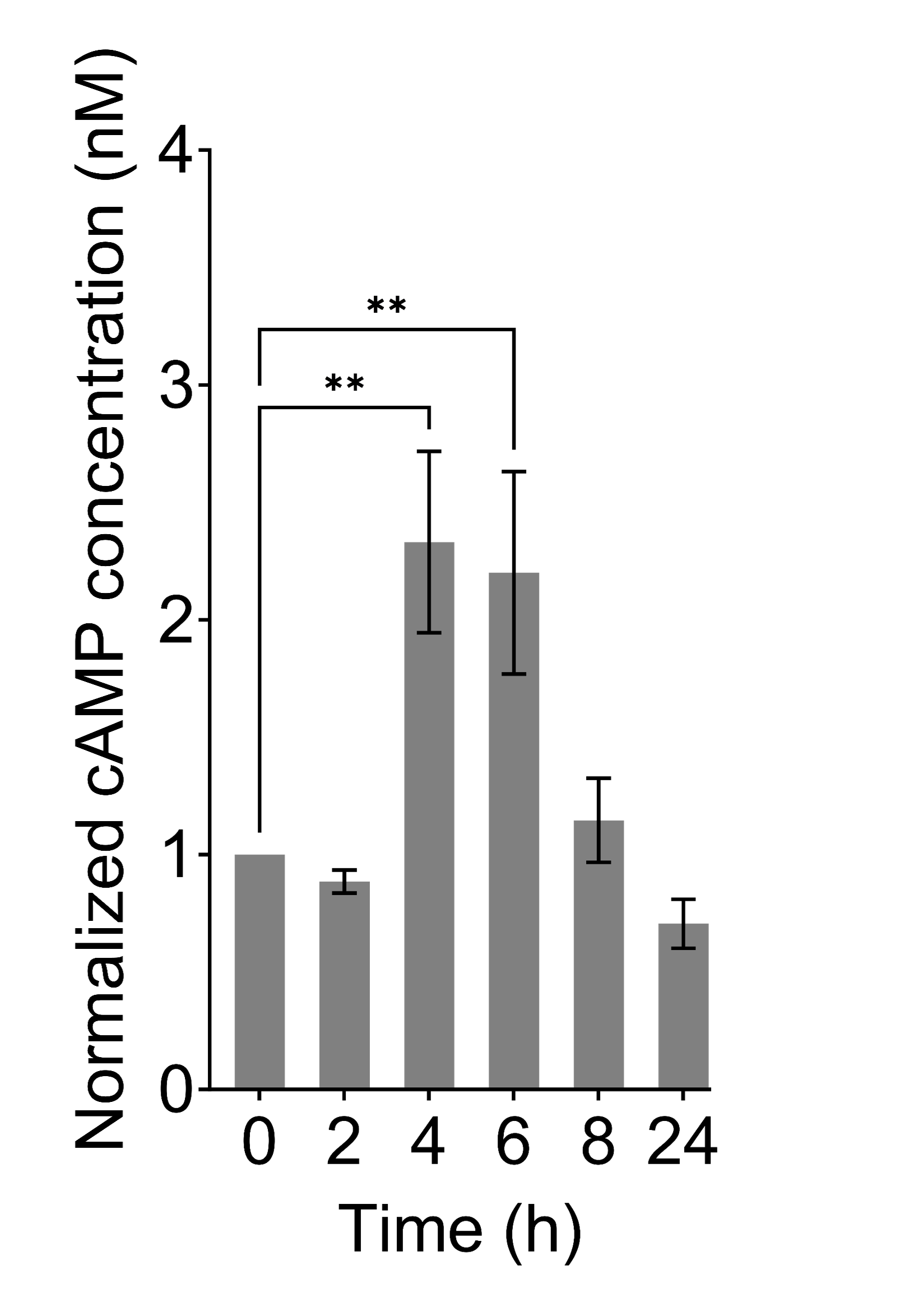

### Figure 1 - figure supplement 5

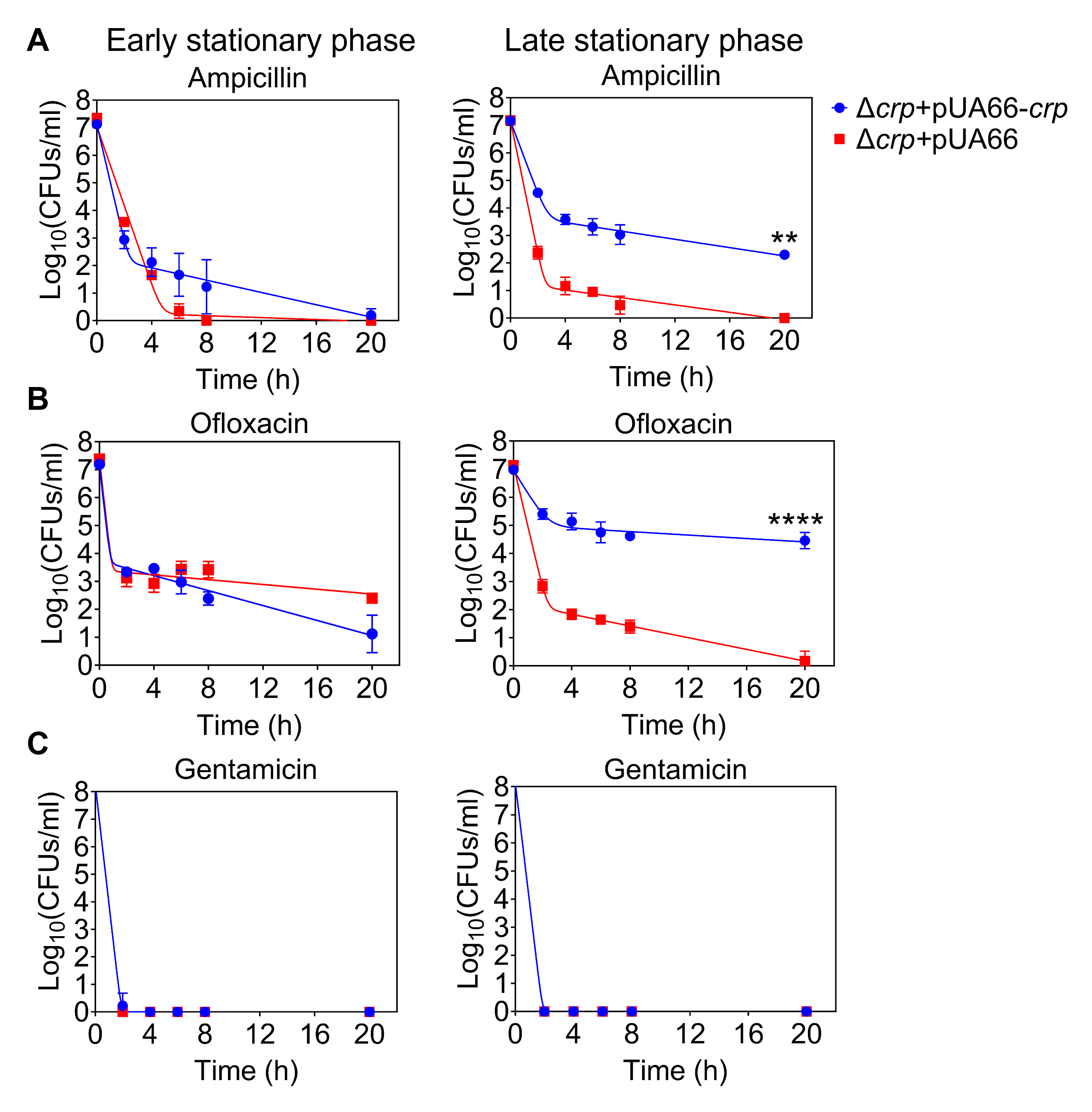

### Figure 1 - figure supplement 6

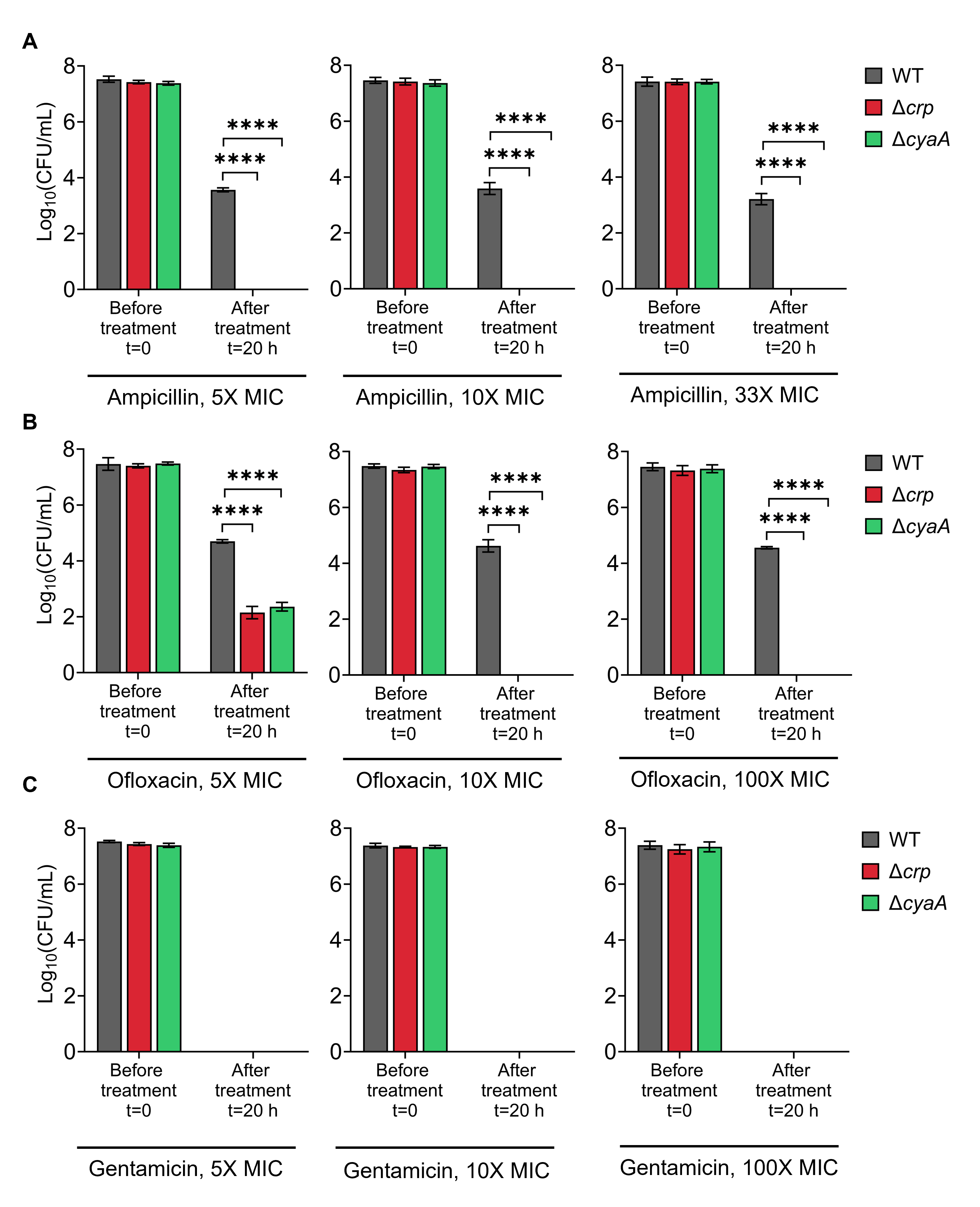

### Figure 1 - figure supplement 7

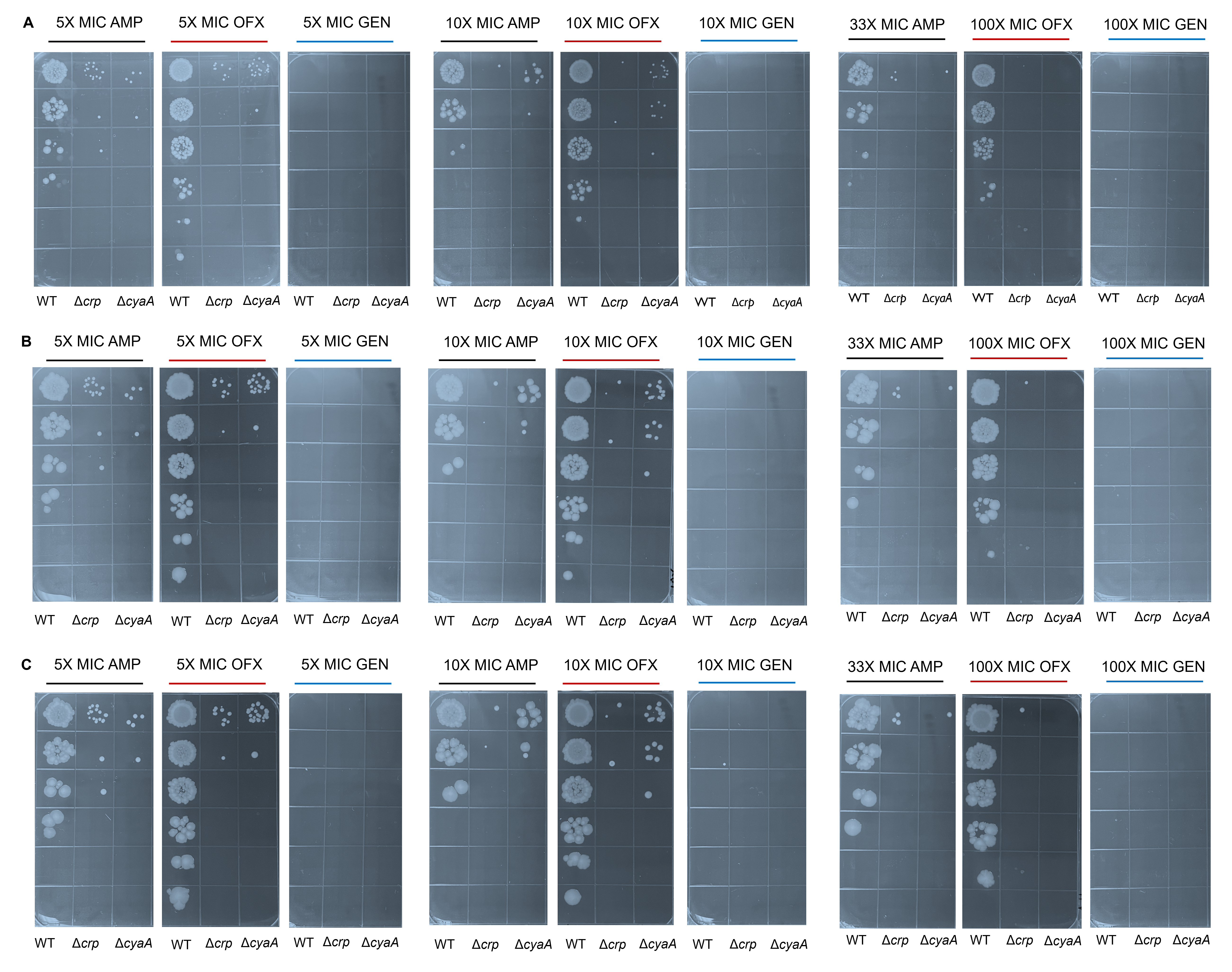

### Figure 1 - figure supplement 8

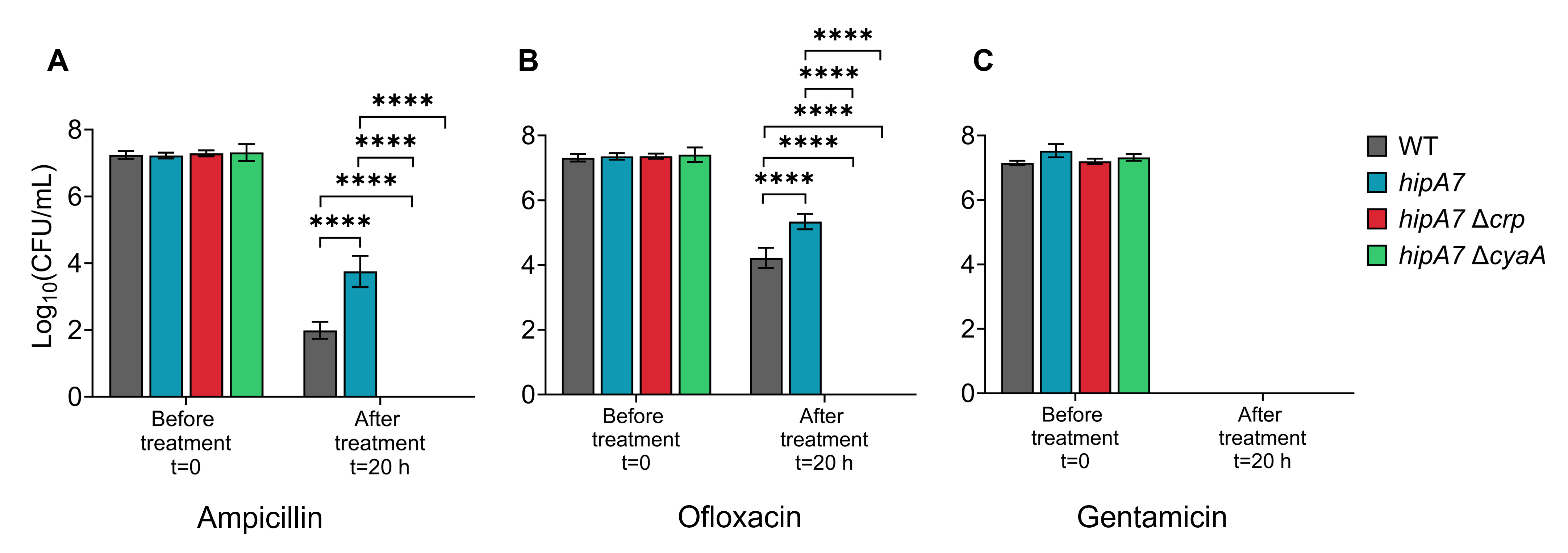

### Figure 2 - figure supplement 1

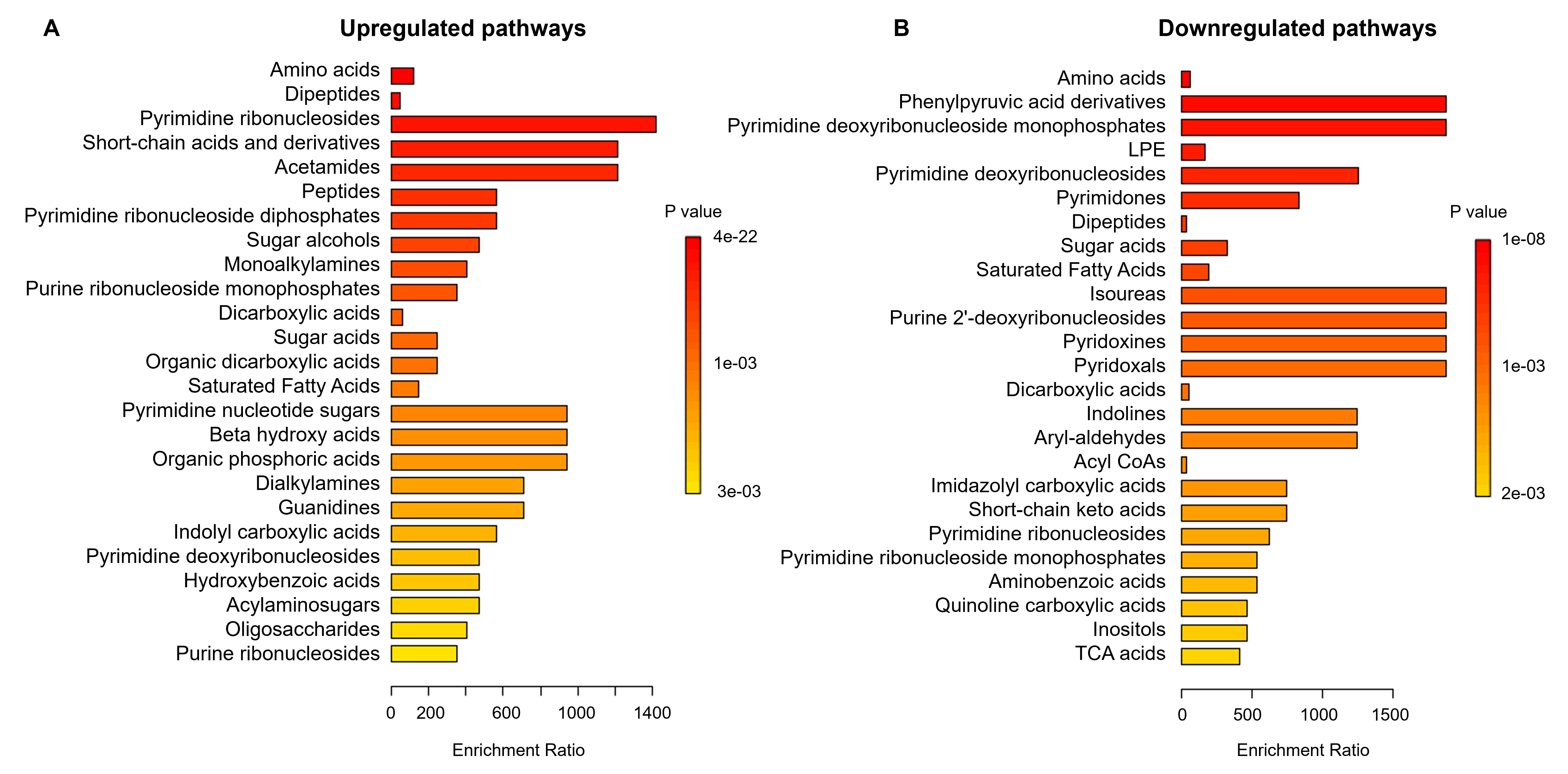

### Figure 2 - figure supplement 2

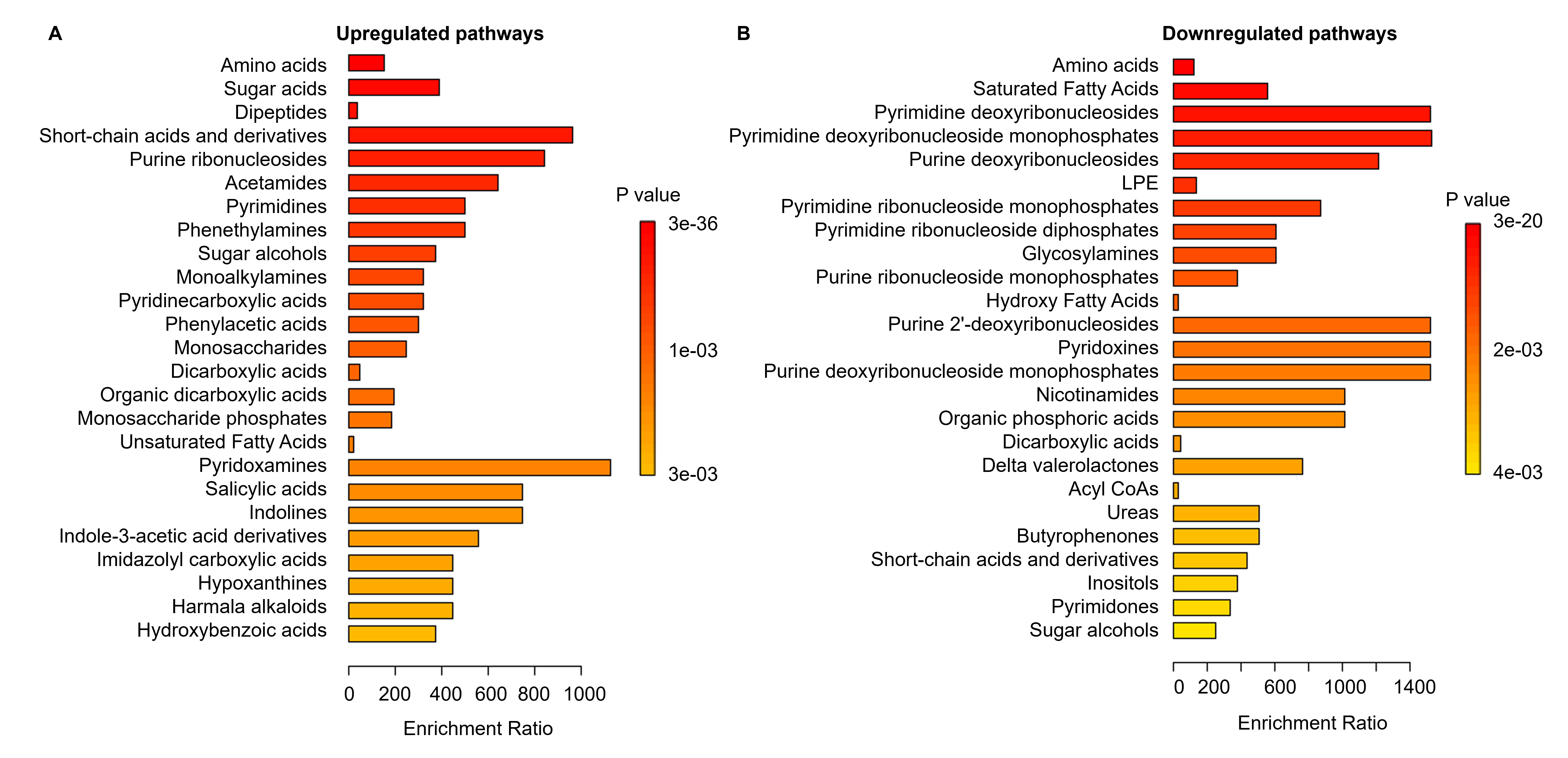

### Figure 2 - figure supplement 3

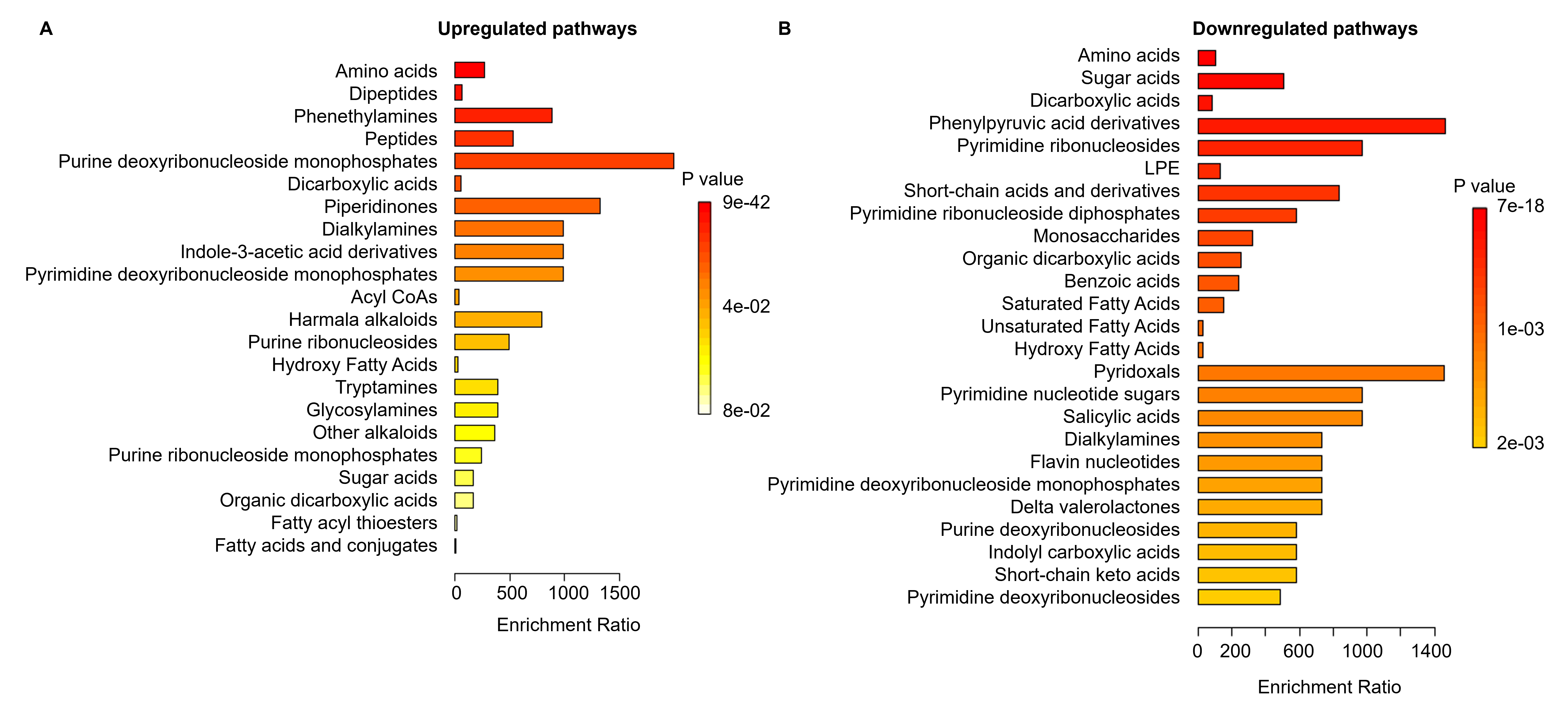

### Figure 3 - figure supplement 1

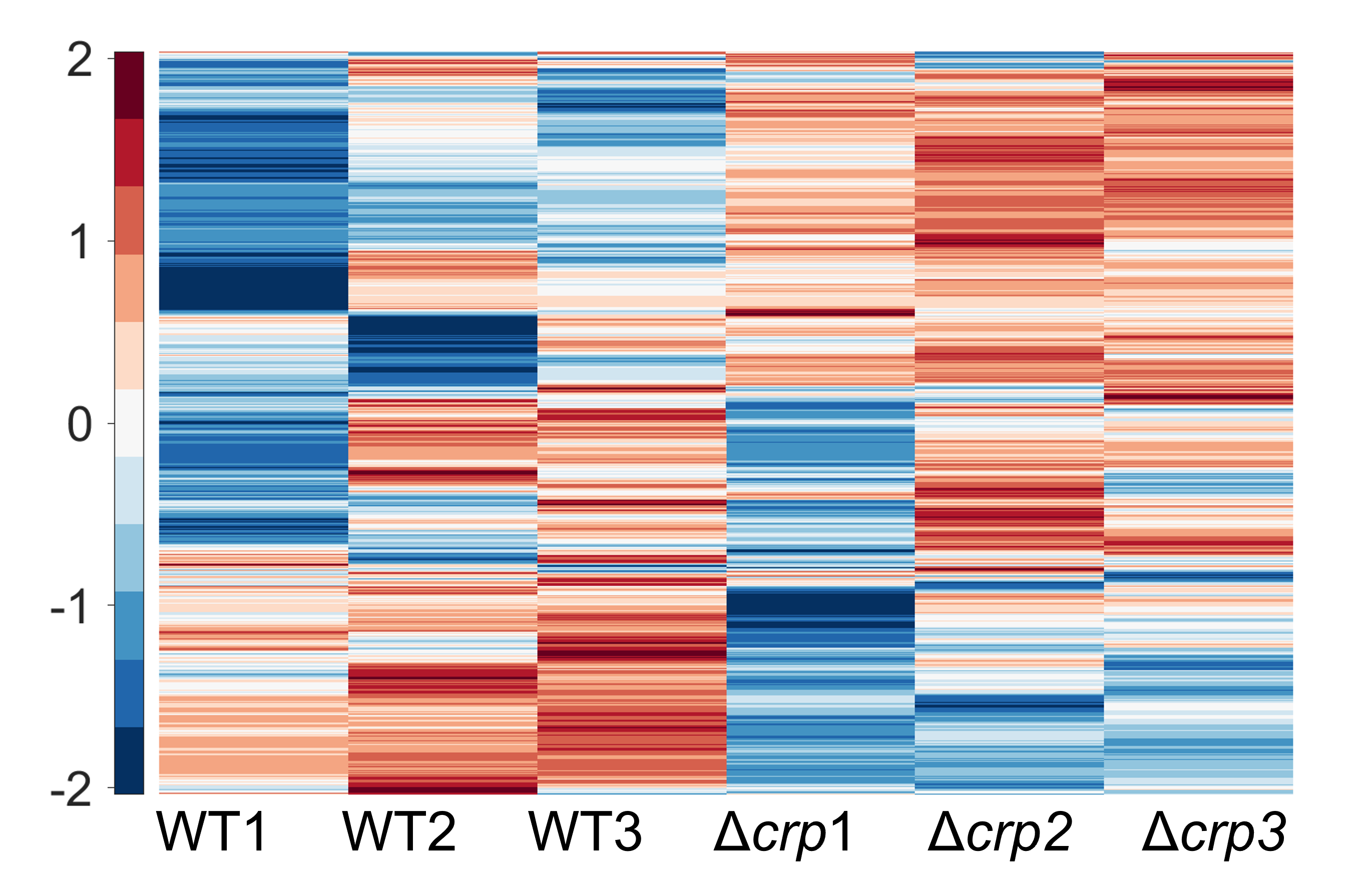

### Figure 4 - figure supplement 1

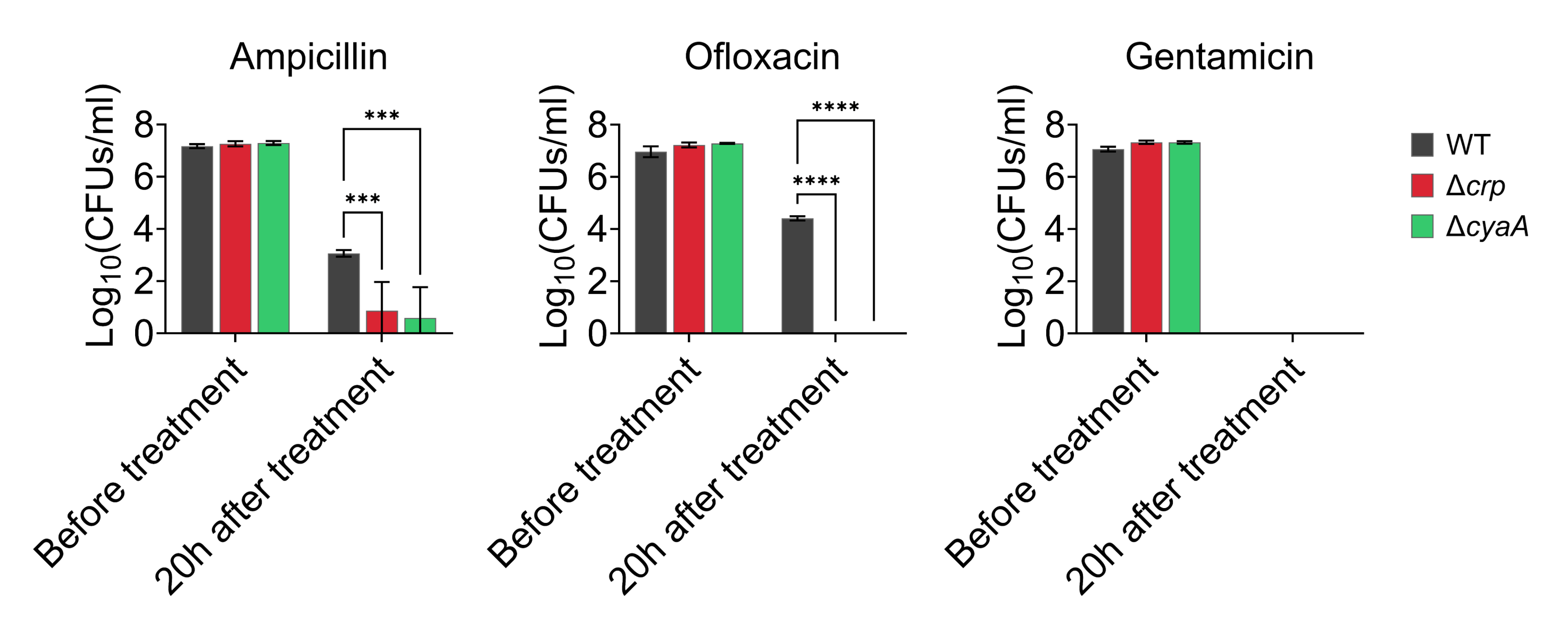

### Figure 4 - figure supplement 2

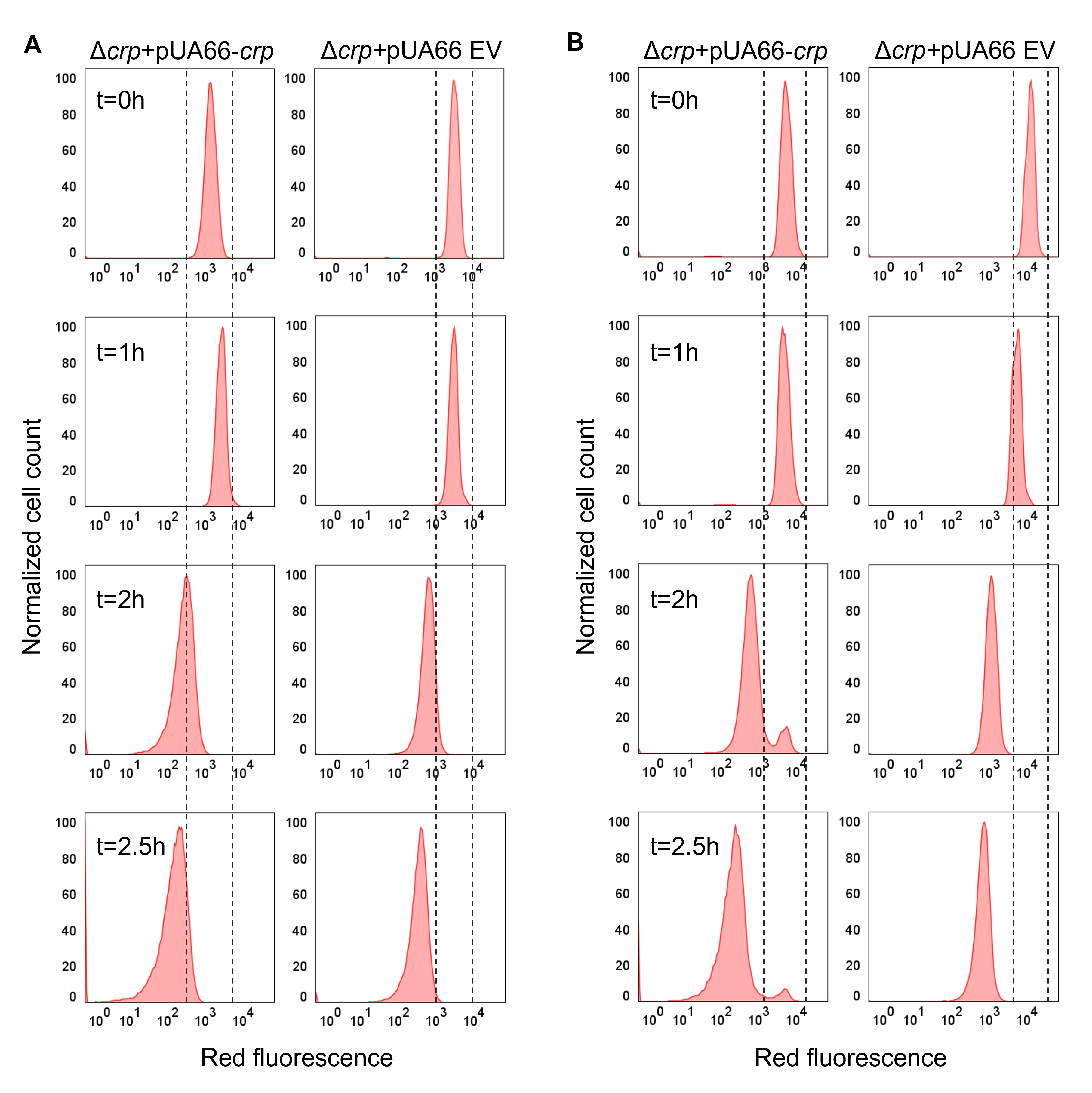

### Figure 5 - figure supplement 1

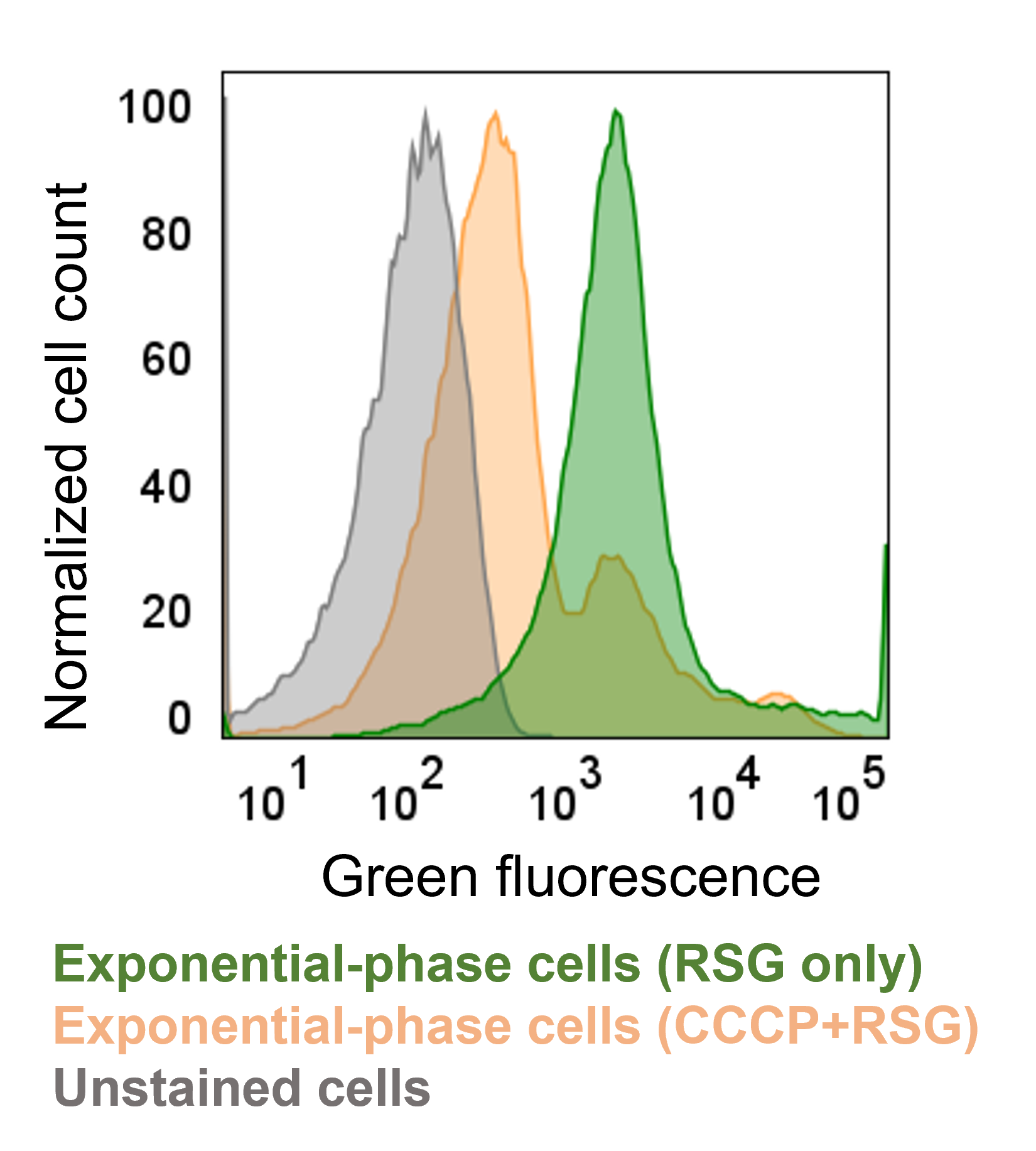

### Figure 5 - figure supplement 2

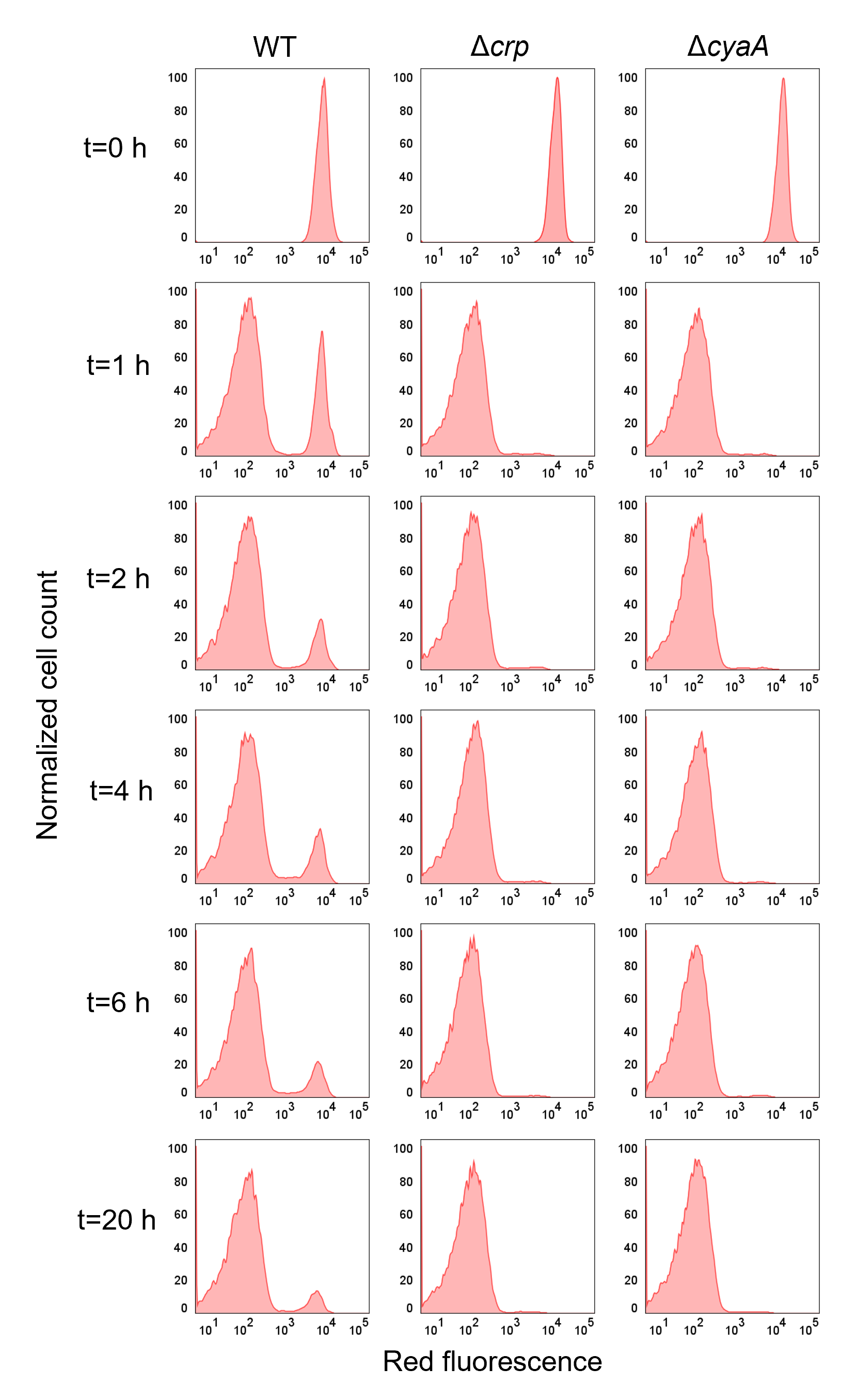

### Figure 5 - figure supplement 3

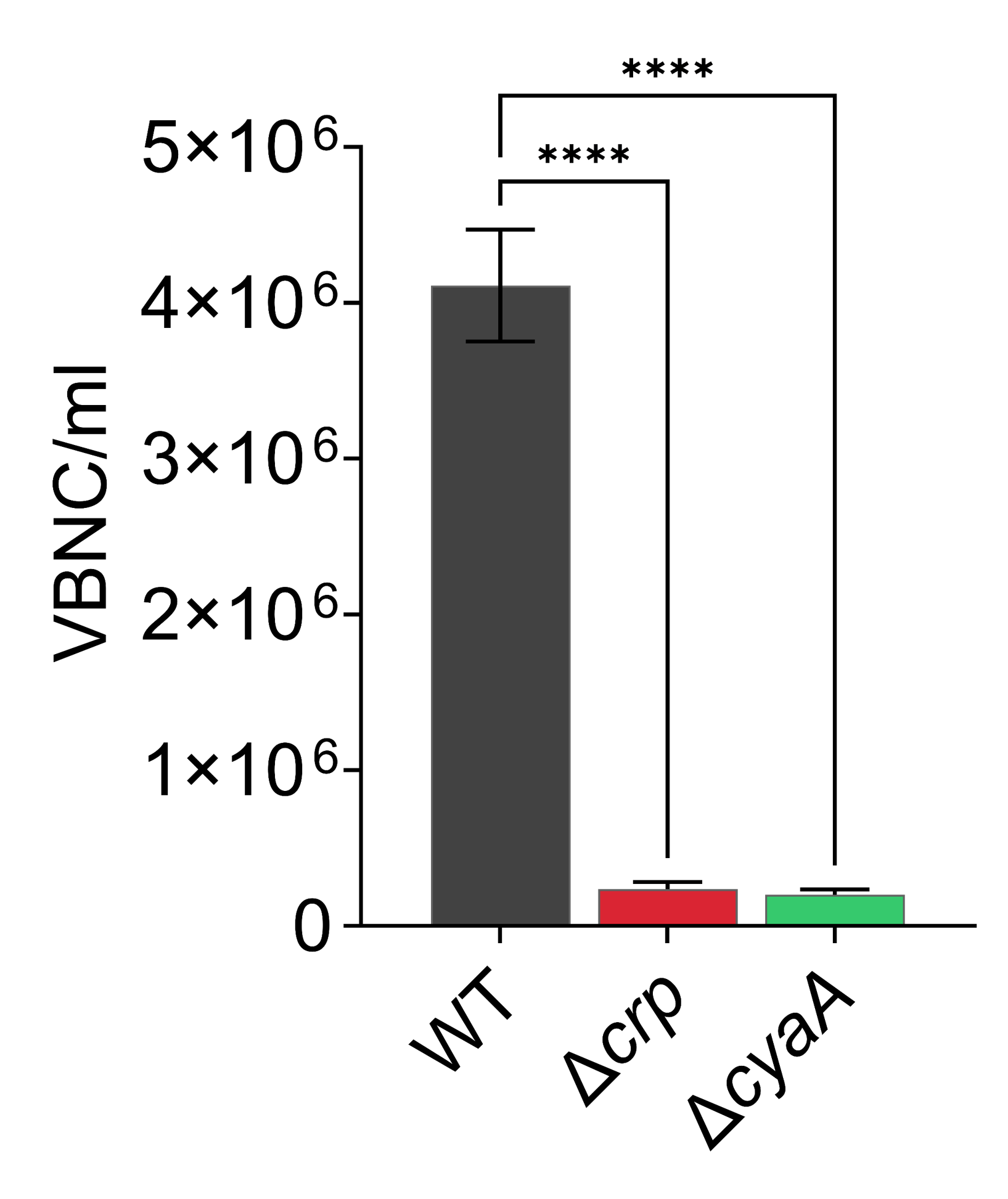

### Figure 5 - figure supplement 4

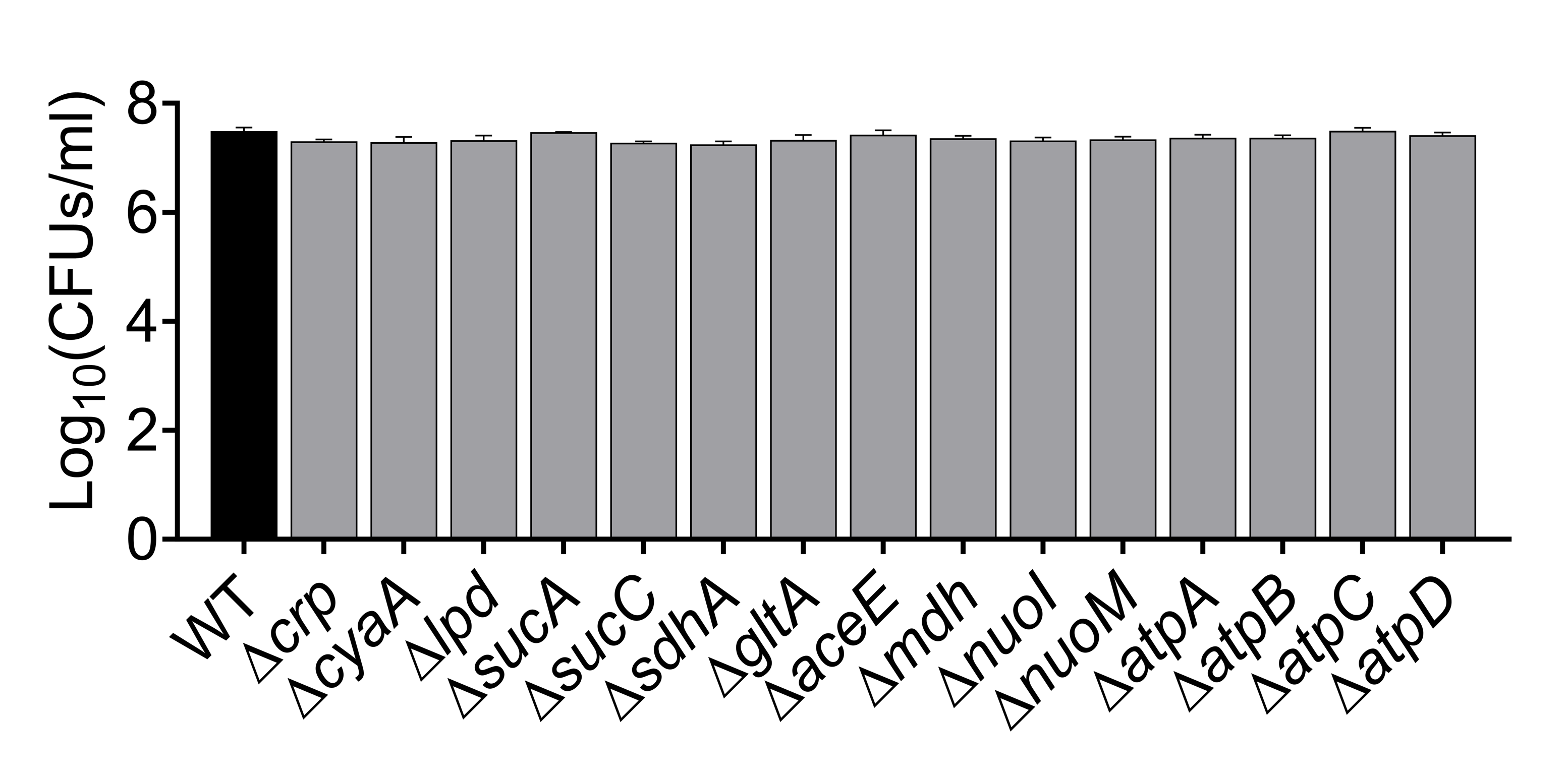
