## Supplementary material for "UNRAVELING CRP/cAMP-MEDIATED METABOLIC REGULATION IN *ESCHERICHIA COLI* PERSISTER CELLS": Key Resources Table

| **Key Resources Table** | | | | |
| --- | --- | --- | --- | --- |
| **Reagent type (species) or resource** | **Designation** | **Source or reference** | **Identifiers** | **Additional information** |
| *Escherichia coli* K-12 MG1655 | Wild Type | Gift from Dr. Mark P. Brynildsen |  |  |
| *Escherichia coli* K-12 MG1655 | *hipA7* | Gift from Dr. Mark P. Brynildsen |  |  |
| *Escherichia coli* K-12 MG1655 | *hipA7*Δ*crp* | This study |  |  |
| *Escherichia coli* K-12 MG1655 | *hipA7*Δ*cyaA* | This study |  |  |
| *Escherichia coli* K-12 BW25113 | Wild Type | Keio collection | Catalog # OEC5042 |  |
| *Escherichia coli* K-12 MG1655 | MO | Gift from Dr. Mark P. Brynildsen |  |  |
| *Escherichia coli* K-12 MG1655 | MO Δ*crp* | This study |  |  |
| *Escherichia coli* K-12 MG1655 | MO Δ*cyaA* | This study |  |  |
| *Escherichia coli* K-12 MG1655 | Δ*crp* | This study |  |  |
| *Escherichia coli* K-12 MG1655 | Δ*cyaA* | This study |  |  |
| *Escherichia coli* K-12 MG1655 | Δ*sucA* | This study |  |  |
| *Escherichia coli* K-12 MG1655 | Δ*lpd* | This study |  |  |
| *Escherichia coli* K-12 MG1655 | Δ*sucC* | This study |  |  |
| *Escherichia coli* K-12 MG1655 | Δ*sdhA* | This study |  |  |
| *Escherichia coli* K-12 MG1655 | Δ*gltA* | This study |  |  |
| *Escherichia coli* K-12 MG1655 | Δ*acnB* | This study |  |  |
| *Escherichia coli* K-12 MG1655 | Δ*aceE* | This study |  |  |
| *Escherichia coli* K-12 MG1655 | Δ*fumA* | This study |  |  |
| *Escherichia coli* K-12 MG1655 | Δ*mdh* | This study |  |  |
| *Escherichia coli* K-12 MG1655 | Δ*nuoI* | This study |  |  |
| *Escherichia coli* K-12 MG1655 | Δ*nuoM* | This study |  |  |
| *Escherichia coli* K-12 MG1655 | Δ*atpA* | This study |  |  |
| *Escherichia coli* K-12 MG1655 | Δ*atpB* | This study |  |  |
| *Escherichia coli* K-12 MG1655 | Δ*atpC* | This study |  |  |
| *Escherichia coli* K-12 MG1655 | Δ*atpD* | This study |  |  |
| *Escherichia coli* K-12 MG1655 | Δ*frdC* | This study |  |  |
| *Escherichia coli* K-12 MG1655 | Δ*pgi* | This study |  |  |
| *Escherichia coli* K-12 MG1655 | Δ*zwf* | This study |  |  |
| *Escherichia coli* K-12 MG1655 | Δ*talA* | This study |  |  |
| *Escherichia coli* K-12 BW25113 | Δ*acnB* | Keio collection | Catalog # OEC4988 |  |
| *Escherichia coli* K-12 BW25113 | Δ*sucA* | Keio collection | Catalog # OEC4988 |  |
| *Escherichia coli* K-12 BW25113 | Δ*sucB* | Keio collection | Catalog # OEC4988 |  |
| *Escherichia coli* K-12 BW25113 | Δ*sucC* | Keio collection | Catalog # OEC4988 |  |
| *Escherichia coli* K-12 BW25113 | Δ*sdhD* | Keio collection | Catalog # OEC4988 |  |
| *Escherichia coli* K-12 BW25113 | Δ*sdhC* | Keio collection | Catalog # OEC4988 |  |
| *Escherichia coli* K-12 BW25113 | Δ*sdhB* | Keio collection | Catalog # OEC4988 |  |
| *Escherichia coli* K-12 BW25113 | Δ*sdhA* | Keio collection | Catalog # OEC4988 |  |
| *Escherichia coli* K-12 BW25113 | Δ*aceF* | Keio collection | Catalog # OEC4988 |  |
| *Escherichia coli* K-12 BW25113 | Δ*talB* | Keio collection | Catalog # OEC4988 |  |
| *Escherichia coli* K-12 BW25113 | Δ*cyoA* | Keio collection | Catalog # OEC4988 |  |
| *Escherichia coli* K-12 BW25113 | Δ*cyoB* | Keio collection | Catalog # OEC4988 |  |
| *Escherichia coli* K-12 BW25113 | Δ*cyoC* | Keio collection | Catalog # OEC4988 |  |
| *Escherichia coli* K-12 BW25113 | Δ*cyoD* | Keio collection | Catalog # OEC4988 |  |
| *Escherichia coli* K-12 BW25113 | Δ*acnA* | Keio collection | Catalog # OEC4988 |  |
| *Escherichia coli* K-12 BW25113 | Δ*icd* | Keio collection | Catalog # OEC4988 |  |
| *Escherichia coli* K-12 BW25113 | Δ*fumA* | Keio collection | Catalog # OEC4988 |  |
| *Escherichia coli* K-12 BW25113 | Δ*fumB* | Keio collection | Catalog # OEC4988 |  |
| *Escherichia coli* K-12 BW25113 | Δ*fumC* | Keio collection | Catalog # OEC4988 |  |
| *Escherichia coli* K-12 BW25113 | Δ*mdh* | Keio collection | Catalog # OEC4988 |  |
| *Escherichia coli* K-12 BW25113 | Δ*pgi* | Keio collection | Catalog # OEC4988 |  |
| *Escherichia coli* K-12 BW25113 | Δ*pfkA* | Keio collection | Catalog # OEC4988 |  |
| *Escherichia coli* K-12 BW25113 | Δ*tpiA* | Keio collection | Catalog # OEC4988 |  |
| *Escherichia coli* K-12 BW25113 | Δ*gpmM* | Keio collection | Catalog # OEC4988 |  |
| *Escherichia coli* K-12 BW25113 | Δ*ppsA* | Keio collection | Catalog # OEC4988 |  |
| *Escherichia coli* K-12 BW25113 | Δ*pykF* | Keio collection | Catalog # OEC4988 |  |
| *Escherichia coli* K-12 BW25113 | Δ*pykA* | Keio collection | Catalog # OEC4988 |  |
| *Escherichia coli* K-12 BW25113 | Δ*maeB* | Keio collection | Catalog # OEC4988 |  |
| *Escherichia coli* K-12 BW25113 | Δ*pck* | Keio collection | Catalog # OEC4988 |  |
| *Escherichia coli* K-12 BW25113 | Δ*rpiB* | Keio collection | Catalog # OEC4988 |  |
| *Escherichia coli* K-12 BW25113 | Δ*rpe* | Keio collection | Catalog # OEC4988 |  |
| *Escherichia coli* K-12 BW25113 | Δ*tktB* | Keio collection | Catalog # OEC4988 |  |
| *Escherichia coli* K-12 BW25113 | Δ*talA* | Keio collection | Catalog # OEC4988 |  |
| *Escherichia coli* K-12 BW25113 | Δ*zwf* | Keio collection | Catalog # OEC4988 |  |
| *Escherichia coli* K-12 BW25113 | Δ*gnd* | Keio collection | Catalog # OEC4988 |  |
| *Escherichia coli* K-12 BW25113 | Δ*ppc* | Keio collection | Catalog # OEC4988 |  |
| *Escherichia coli* K-12 BW25113 | Δ*frdA* | Keio collection | Catalog # OEC4988 |  |
| *Escherichia coli* K-12 BW25113 | Δ*frdB* | Keio collection | Catalog # OEC4988 |  |
| *Escherichia coli* K-12 BW25113 | Δ*frdC* | Keio collection | Catalog # OEC4988 |  |
| *Escherichia coli* K-12 BW25113 | Δ*frdD* | Keio collection | Catalog # OEC4988 |  |
| *Escherichia coli* K-12 BW25113 | Δ*adhE* | Keio collection | Catalog # OEC4988 |  |
| *Escherichia coli* K-12 BW25113 | Δ*pflB* | Keio collection | Catalog # OEC4988 |  |
| *Escherichia coli* K-12 BW25113 | Δ*aceB* | Keio collection | Catalog # OEC4988 |  |
| *Escherichia coli* K-12 BW25113 | Δ*aceA* | Keio collection | Catalog # OEC4988 |  |
| *Escherichia coli* K-12 BW25113 | Δ*glcB* | Keio collection | Catalog # OEC4988 |  |
| *Escherichia coli* K-12 BW25113 | Δ*nuoA* | Keio collection | Catalog # OEC4988 |  |
| *Escherichia coli* K-12 BW25113 | Δ*nuoH* | Keio collection | Catalog # OEC4988 |  |
| *Escherichia coli* K-12 BW25113 | Δ*nuoJ* | Keio collection | Catalog # OEC4988 |  |
| *Escherichia coli* K-12 BW25113 | Δ*nuoK* | Keio collection | Catalog # OEC4988 |  |
| *Escherichia coli* K-12 BW25113 | Δ*nuoL* | Keio collection | Catalog # OEC4988 |  |
| *Escherichia coli* K-12 BW25113 | Δ*nuoM* | Keio collection | Catalog # OEC4988 |  |
| *Escherichia coli* K-12 BW25113 | Δ*nuoN* | Keio collection | Catalog # OEC4988 |  |
| *Escherichia coli* K-12 BW25113 | Δ*nuoB* | Keio collection | Catalog # OEC4988 |  |
| *Escherichia coli* K-12 BW25113 | Δ*nuoE* | Keio collection | Catalog # OEC4988 |  |
| *Escherichia coli* K-12 BW25113 | Δ*nuoF* | Keio collection | Catalog # OEC4988 |  |
| *Escherichia coli* K-12 BW25113 | Δ*nuoG* | Keio collection | Catalog # OEC4988 |  |
| *Escherichia coli* K-12 BW25113 | Δ*nuoI* | Keio collection | Catalog # OEC4988 |  |
| *Escherichia coli* K-12 BW25113 | Δ*appC* | Keio collection | Catalog # OEC4988 |  |
| *Escherichia coli* K-12 BW25113 | Δ*appB* | Keio collection | Catalog # OEC4988 |  |
| *Escherichia coli* K-12 BW25113 | Δ*gltA* | Keio collection | Catalog # OEC4988 |  |
| *Escherichia coli* K-12 BW25113 | Δ*lpd* | Keio collection | Catalog # OEC4988 |  |
| *Escherichia coli* K-12 BW25113 | Δ*fbp* | Keio collection | Catalog # OEC4988 |  |
| *Escherichia coli* K-12 BW25113 | Δ*aceE* | Keio collection | Catalog # OEC4988 |  |
| *Escherichia coli* K-12 BW25113 | Δ*tktA* | Keio collection | Catalog # OEC4988 |  |
| *Escherichia coli* K-12 BW25113 | Δ*pta* | Keio collection | Catalog # OEC4988 |  |
| *Escherichia coli* K-12 BW25113 | Δ*ackA* | Keio collection | Catalog # OEC4988 |  |
| *Escherichia coli* K-12 BW25113 | Δ*ldhA* | Keio collection | Catalog # OEC4988 |  |
| *Escherichia coli* K-12 BW25113 | Δ*dld* | Keio collection | Catalog # OEC4988 |  |
| *Escherichia coli* K-12 BW25113 | Δ*cydB* | Keio collection | Catalog # OEC4988 |  |
| *Escherichia coli* K-12 BW25113 | Δ*poxB* | Keio collection | Catalog # OEC4988 |  |
| *Escherichia coli* K-12 BW25113 | Δ*glpX* | Keio collection | Catalog # OEC4988 |  |
| *Escherichia coli* K-12 BW25113 | Δ*ybhA* | Keio collection | Catalog # OEC4988 |  |
| *Escherichia coli* K-12 BW25113 | Δ*cydX* | Keio collection | Catalog # OEC4988 |  |
| *Escherichia coli* K-12 BW25113 | Δ*fumE* | Keio collection | Catalog # OEC4988 |  |
| *Escherichia coli* K-12 BW25113 | Δ*yggF* | Keio collection | Catalog # OEC4988 |  |
| *Escherichia coli* K-12 BW25113 | Δ*ccp* | Keio collection | Catalog # OEC4988 |  |
| *Escherichia coli* K-12 BW25113 | Δ*yieF* | Keio collection | Catalog # OEC4988 |  |
| *Escherichia coli* K-12 BW25113 | Δ*wrbA* | Keio collection | Catalog # OEC4988 |  |
| *Escherichia coli* K-12 BW25113 | Δ*atpC* | Keio collection | Catalog # OEC4988 |  |
| *Escherichia coli* K-12 BW25113 | Δ*atpD* | Keio collection | Catalog # OEC4988 |  |
| *Escherichia coli* K-12 BW25113 | Δ*fdhF* | Keio collection | Catalog # OEC4988 |  |
| *Escherichia coli* K-12 BW25113 | Δ*nrfD* | Keio collection | Catalog # OEC4988 |  |
| *Escherichia coli* K-12 BW25113 | Δ*nrfC* | Keio collection | Catalog # OEC4988 |  |
| *Escherichia coli* K-12 BW25113 | Δ*nrfA* | Keio collection | Catalog # OEC4988 |  |
| *Escherichia coli* K-12 BW25113 | Δ*putA* | Keio collection | Catalog # OEC4988 |  |
| *Escherichia coli* K-12 BW25113 | Δ*dmsC* | Keio collection | Catalog # OEC4988 |  |
| *Escherichia coli* K-12 BW25113 | Δ*torC* | Keio collection | Catalog # OEC4988 |  |
| *Escherichia coli* K-12 BW25113 | Δ*torA* | Keio collection | Catalog # OEC4988 |  |
| *Escherichia coli* K-12 BW25113 | Δ*hyaA* | Keio collection | Catalog # OEC4988 |  |
| *Escherichia coli* K-12 BW25113 | Δ*hyaB* | Keio collection | Catalog # OEC4988 |  |
| *Escherichia coli* K-12 BW25113 | Δ*hyaC* | Keio collection | Catalog # OEC4988 |  |
| *Escherichia coli* K-12 BW25113 | Δ*kefF* | Keio collection | Catalog # OEC4988 |  |
| *Escherichia coli* K-12 BW25113 | Δ*narV* | Keio collection | Catalog # OEC4988 |  |
| *Escherichia coli* K-12 BW25113 | Δ*narI* | Keio collection | Catalog # OEC4988 |  |
| *Escherichia coli* K-12 BW25113 | Δ*kduI* | Keio collection | Catalog # OEC4988 |  |
| *Escherichia coli* K-12 BW25113 | Δ*eutE* | Keio collection | Catalog # OEC4988 |  |
| *Escherichia coli* K-12 BW25113 | Δ*hycE* | Keio collection | Catalog # OEC4988 |  |
| *Escherichia coli* K-12 BW25113 | Δ*hycG* | Keio collection | Catalog # OEC4988 |  |
| *Escherichia coli* K-12 BW25113 | Δ*edd* | Keio collection | Catalog # OEC4988 |  |
| *Escherichia coli* K-12 BW25113 | Δ*eda* | Keio collection | Catalog # OEC4988 |  |
| *Escherichia coli* K-12 BW25113 | Δ*fdnG* | Keio collection | Catalog # OEC4988 |  |
| *Escherichia coli* K-12 BW25113 | Δ*fdnI* | Keio collection | Catalog # OEC4988 |  |
| *Escherichia coli* K-12 BW25113 | Δ*glpD* | Keio collection | Catalog # OEC4988 |  |
| *Escherichia coli* K-12 BW25113 | Δ*glpA* | Keio collection | Catalog # OEC4988 |  |
| *Escherichia coli* K-12 BW25113 | Δ*glpB* | Keio collection | Catalog # OEC4988 |  |
| *Escherichia coli* K-12 BW25113 | Δ*glpC* | Keio collection | Catalog # OEC4988 |  |
| *Escherichia coli* K-12 BW25113 | Δ*hybO* | Keio collection | Catalog # OEC4988 |  |
| *Escherichia coli* K-12 BW25113 | Δ*hybC* | Keio collection | Catalog # OEC4988 |  |
| *Escherichia coli* K-12 BW25113 | Δ*narY* | Keio collection | Catalog # OEC4988 |  |
| *Escherichia coli* K-12 BW25113 | Δ*narZ* | Keio collection | Catalog # OEC4988 |  |
| *Escherichia coli* K-12 BW25113 | Δ*napG* | Keio collection | Catalog # OEC4988 |  |
| *Escherichia coli* K-12 BW25113 | Δ*napH* | Keio collection | Catalog # OEC4988 |  |
| *Escherichia coli* K-12 BW25113 | Δ*napA* | Keio collection | Catalog # OEC4988 |  |
| *Escherichia coli* K-12 BW25113 | Δ*atpA* | Keio collection | Catalog # OEC4988 |  |
| *Escherichia coli* K-12 BW25113 | Δ*purT* | Keio collection | Catalog # OEC4988 |  |
| *Escherichia coli* K-12 BW25113 | Δ*fdoG* | Keio collection | Catalog # OEC4988 |  |
| *Escherichia coli* K-12 BW25113 | Δ*fdoI* | Keio collection | Catalog # OEC4988 |  |
| *Escherichia coli* K-12 BW25113 | Δ*atpB* | Keio collection | Catalog # OEC4988 |  |
| *Escherichia coli* K-12 BW25113 | Δ*atpE* | Keio collection | Catalog # OEC4988 |  |
| *Escherichia coli* K-12 BW25113 | Δ*atpF* | Keio collection | Catalog # OEC4988 |  |
| *Escherichia coli* K-12 BW25113 | Δ*atpH* | Keio collection | Catalog # OEC4988 |  |
| *Escherichia coli* K-12 BW25113 | Δ*phoA* | Keio collection | Catalog # OEC4988 |  |
| *Escherichia coli* K-12 BW25113 | Δ*adhP* | Keio collection | Catalog # OEC4988 |  |
| *Escherichia coli* K-12 BW25113 | Δ*eutD* | Keio collection | Catalog # OEC4988 |  |
| *Escherichia coli* K-12 BW25113 | Δ*dmsA* | Keio collection | Catalog # OEC4988 |  |
| *Escherichia coli* K-12 BW25113 | Δ*hybB* | Keio collection | Catalog # OEC4988 |  |
| *Escherichia coli* K-12 BW25113 | Δ*napB* | Keio collection | Catalog # OEC4988 |  |
| *Escherichia coli* K-12 BW25113 | Δ*atpI* | Keio collection | Catalog # OEC4988 |  |
| *Escherichia coli* K-12 BW25113 | Δ*maeA* | Keio collection | Catalog # OEC4988 |  |
| *Escherichia coli* K-12 BW25113 | Δ*pfkB* | Keio collection | Catalog # OEC4988 |  |
| *Escherichia coli* K-12 BW25113 | Δ*fbaB* | Keio collection | Catalog # OEC4988 |  |
| *Escherichia coli* K-12 BW25113 | Δ*rpiA* | Keio collection | Catalog # OEC4988 |  |
| *Escherichia coli* K-12 BW25113 | Δ*fumD* | Keio collection | Catalog # OEC4988 |  |
| *Escherichia coli* K-12 BW25113 | Δ*sucD* | Keio collection | Catalog # OEC4988 |  |
| *Escherichia coli* K-12 BW25113 | Δ*fdnH* | Keio collection | Catalog # OEC4988 |  |
| *Escherichia coli* K-12 BW25113 | Δ*dmsB* | Keio collection | Catalog # OEC4988 |  |
| *Escherichia coli* K-12 BW25113 | Δ*ndh* | Keio collection | Catalog # OEC4988 |  |
| *Escherichia coli* K-12 BW25113 | Δ*narG* | Keio collection | Catalog # OEC4988 |  |
| *Escherichia coli* K-12 BW25113 | Δ*narH* | Keio collection | Catalog # OEC4988 |  |
| *Escherichia coli* K-12 BW25113 | Δ*tdcE* | Keio collection | Catalog # OEC4988 |  |
| *Escherichia coli* K-12 BW25113 | Δ*hycB* | Keio collection | Catalog # OEC4988 |  |
| *Escherichia coli* K-12 BW25113 | Δ*hycC* | Keio collection | Catalog # OEC4988 |  |
| *Escherichia coli* K-12 BW25113 | Δ*hycD* | Keio collection | Catalog # OEC4988 |  |
| *Escherichia coli* K-12 BW25113 | Δ*fdoH* | Keio collection | Catalog # OEC4988 |  |
| *Escherichia coli* K-12 BW25113 | Δ*hybA* | Keio collection | Catalog # OEC4988 |  |
| *Escherichia coli* K-12 BW25113 | Δ*nuoC* | Keio collection | Catalog # OEC4988 |  |
| *Escherichia coli* K-12 BW25113 | Δ*atpG* | Keio collection | Catalog # OEC4988 |  |
| Bacterial Plasmids | *pMSs201 (kan^R^)* | Dharmacon Promoter Library | Catalog# OEC4988 |  |
| Bacterial Plasmids | pUA66-EV (empty vector) | Gift from Dr. Mark P. Brynildsen |  | A DNA fragment including *T5* promoter, *Kan^R^* gene, pUA66 origin of replication and *lacI^q^* was amplified from the pUA66-*gfp* plasmid with primers having BspHI cut sites. The amplified DNA fragment was digested with BspHI, and then self-ligated to obtain the modified pUA66-EV that does not have the *gfp* gene. |
| Bacterial Plasmids | pUA66-*crp* | This study |  | The *crp* gene with its promoter was amplified from the genomic DNA of *E.coli*, using forward and reverse primers with BglII and ScaI restriction enzyme cut sites, respectively. The pUA66-*gfp* plasmid was double digested with BglII and ScaI to remove T5 promoter region and *gfp* gene. Then, the digested *crp* gene with its promoter and plasmid were ligated to generate pUA66-*crp*. |
| *Escherichia coli* K-12 MG1655 | pMSs201 P*_sdhABCD_*-*gfp* | This study |  |  |
| *Escherichia coli* K-12 MG1655 | Δ*crp* pMSs201 P*_sdhABCD_*-*gfp* | This study |  |  |
| *Escherichia coli* K-12 MG1655 | Δ*cyaA* pMSs201 P*_sdhABCD_*-*gfp* | This study |  |  |
| *Escherichia coli* K-12 MG1655 | pMSs201 P*_cyaA_*-*gfp* | This study |  |  |
| *Escherichia coli* K-12 MG1655 | Δ*crp* pMSs201 P*_cyaA_*-*gfp* | This study |  |  |
| *Escherichia coli* K-12 MG1655 | Δ*cyaA* pMSs201 P*_cyaA_*-*gfp* | This study |  |  |
| *Escherichia coli* K-12 MG1655 | Δ*crp* pUA66-EV | This study |  |  |
| *Escherichia coli* K-12 MG1655 | Δ*crp* pUA66-*crp* | This study |  |  |
| Software, algorithm | Prism (version 10.3.0) | GraphPad | RRID: SCR_002798 | http://www.graphpad.com/ |
| Software, algorithm | FlowJo  (version 10.8.1) | Becton, Dickinson & Company | RRID: SCR_008520 | https://www.flowjo.com/ |
| Software, algorithm | MATLAB  (version R2020b) | MathWorks | RRID:  SCR_001622 | https://www.mathworks.com/ |

| **Oligonucleotides for the construction and verification of mutant strains.** | | | | | | |  |  |  | |
| --- | --- | --- | --- | --- | --- | --- | --- | --- | --- | --- |
| **Oligonucleotides to Generate Gene Deletions** | | | |  |  |  |  |  |  |  |
| **Mutation** | **Forward Primer (5’ to 3’)** | **Reverse Primer (5’ to 3’)** | **Source** |  | | | | | |  |
| Δ*crp*::KAN(R) | TCTGGCTCTGGAGAAAGCTTATAACAGAGGATAACCGCGCGTGTAGGCTGGAGCTGCTTC | AAAATGGCGCGCTACCAGGTAACGCGCCACTCCGACGGGATTAACGGCTGACATGGGAAT | Integrated DNA Technologies, Inc. |  | | | | | |  |
| Δ*cyaA*::KAN(R) | GAATCACAGTCATGACGGGTAGCAAATCAGGCGATACGTCGTGTAGGCTGGAGCTGCTTC | AGATTGCATGCCGGATAAGCCTCGCTTTCCGGCACGTTCATTAACGGCTGACATGGGAAT | Integrated DNA Technologies, Inc. |  | | | | | |  |
| Δ*sucA*::KAN(R) | ACGGCGAAGTAAGCATAAAAAAGATGCTTAAGGGATCACGGTGTAGGCTGGAGCTGCTTC | GGTCAGGGACCAGAATATCTACGCTACTCATTGTGTATCCTTTATTTAACGGCTGACATGGGAAT | Integrated DNA Technologies, Inc. |  | | | | | |  |
| Δ*lpd*::KAN(R) | GACGGGTATGACCGCCGGAGATAAATATATAGAGGTCATGGTGTAGGCTGGAGCTGCTTC | GCCGCTTTTTTAATTGCCGGATGTTCCGGCAAACGAAAAATTAACGGCTGACATGGGAAT | Integrated DNA Technologies, Inc. |  | | | | | |  |
| Δ*sucC*::KAN(R) | GGTTTAAAAGATAACGATTACTGAAGGATGGACAGAACACGTGTAGGCTGGAGCTGCTTC | TGGCAGATAACCTTGGTGTTTTTATCGATTAAAATGGACATTAACGGCTGACATGGGAAT | Integrated DNA Technologies, Inc. |  | | | | | |  |
| Δ*sdhA*::KAN(R) | TTTACGTGATTTATG GATTCGTTGTGGTGT GGGGTGTGTGGTGTA GGCTGGAGCTGCTTC | GATAAATTGAAAACT CGAGTCTCATTTTCC TGTCTCCGCATTAAC GGCTGACATGGGAAT | Integrated DNA Technologies, Inc. |  | | | | | |  |
| Δ*gltA*::KAN(R) | TAAGTTCCGGCAGTCTTACGCAATAAGGCGCTAAGGAGACCTTAAGTGTAGGCTGGAGCTGCTTC | CCCGCCATATGAACGGCGGGTTAAAATATTTACAACTTAGCAATCAACCATTAACGGCTGACATGGGAAT | Integrated DNA Technologies, Inc. |  | | | | | |  |
| Δ*acnB*::KAN(R) | AATCGCCTGCCGCACTATGACAATGAGAGCGAGGAGAACCGTCGTGTAGGCTGGAGCTGCTTC | GGGCATTGTGTCGTTTATGCGCAGCGCGTGCGCTGACTTTTTAACGGCTGACATGGGAAT | Integrated DNA Technologies, Inc. |  | | | | | |  |
| Δ*aceE*::KAN(R) | GGTTCCAGAAAACTCAACGTTATTAGATAGATAAGGAATAACCCGTGTAGGCTGGAGCTGCTTC | GCCCCGATGTCCGGTACTTTGATTTCGATAGCCATTATTCTTTTACCTCTTAACGGCTGACATGGGAAT | Integrated DNA Technologies, Inc. |  | | | | | |  |
| Δ*fumA*::KAN(R) | GCCCAGAGCATAACCAAACCAGGCAGTAAGTGAGAGAACAGTGTAGGCTGGAGCTGCTTC | TCGTGCCATGTAAAAAAACCGCCCCGAAGGGCGGCTCTGTTTAACGGCTGACATGGGAAT | Integrated DNA Technologies, Inc. |  | | | | | |  |
| Δ*mdh*::KAN(R) | GCGGAGCAACATATCTTAGTTTATCAATATAATAAGGAGTTTAGGGTGTAGGCTGGAGCTGCTTC | CCGGAGTCTGTGCTCCGGTTTTTTATTATCCGCTAATCAATTAACGGCTGACATGGGAAT | Integrated DNA Technologies, Inc. |  | | | | | |  |
| Δ*nuoI*::KAN(R) | CTGTCATTCTCTGGCAGGCGCAATAAGGGGCAATAAGACCGTGTAGGCTGGAGCTGCTTC | AGGCCACAGATATAAAAAGCGAACTCCATTGCCCCTCTCCTTAACGGCTGACATGGGAAT | Integrated DNA Technologies, Inc. |  | | | | | |  |
| Δ*nuoM*::KAN(R) | TCCGGTCCTGACGGGACTTTTACAAGGAATAAAGATCGCCGTGTAGGCTGGAGCTGCTTC | GCAGTGCGATCAGGTTTTGTGGAGTTATTGTCATGGCGATTTAACGGCTGACATGGGAAT | Integrated DNA Technologies, Inc. |  | | | | | |  |
| Δ*atpA*::KAN(R) | GCGCCTTGCAGACGTCTTGCAGTCTTAAGGGGACTGGAGCGTGTAGGCTGGAGCTGCTTC | TCAATGCCTTGCGGCCTGCCCTAAGGCAAGCCGCCAGACGTTAACGGCTGACATGGGAAT | Integrated DNA Technologies, Inc. |  | | | | | |  |
| Δ*atpB*::KAN(R) | TGGCACCGGCTGTAATTAACAACAAAGGGTAAAAGGCATCGTGTAGGCTGGAGCTGCTTC | CTCCAGTTTGTTTCAGTTAAAACGTAGTAGTGTTGGTAAATTAACGGCTGACATGGGAAT | Integrated DNA Technologies, Inc. |  | | | | | |  |
| Δ*atpC*::KAN(R) | GGAAAAAGCCAAAAAACTTTAACGCCTTAATCGGAGGGTGATGTGTAGGCTGGAGCTGCTTC | GCCTGTTTCCAGACTGGCTTTTGTGCTTTTCAAGCCGGTGTTAACGGCTGACATGGGAAT | Integrated DNA Technologies, Inc. |  | | | | | |  |
| Δ*atpD*::KAN(R) | CCGCCGCGGTTTAAACAGGTTATTTCGTAGAGGATTTAAGGTGTAGGCTGGAGCTGCTTC | AGGTGGTAAGTCATTGCCATATCACCCTCCGATTAAGGCGTTAACGGCTGACATGGGAAT | Integrated DNA Technologies, Inc. |  | | | | | |  |
| Δ*frdC*::KAN(R) | TTTCTTATCGCGACCCTGAAACCACGCTAAGGAGTGCAACGTGTAGGCTGGAGCTGCTTC | GTCAGAACGCTTTGGATTTGGATTAATCATCTCAGGCTCCTTAACGGCTGACATGGGAAT | Integrated DNA Technologies, Inc. |  | | | | | |  |
| Δ*pgi*::KAN(R) | GCTACAATCTTCCAAAGTCACAATTCTCAAAATCAGAAGAGTATTGCTAGTGTAGGCTGGAGCTGCTTC | GCGGCGTGAACGCCTTATCCGGCCTACATATCGACGATGATTAACGGCTGACATGGGAAT | Integrated DNA Technologies, Inc. |  | | | | | |  |
| Δ*zwf*::KAN(R) | CTGGCTTAAGTACCGGGTTAGTTAACTTAAGGAGAATGACGTGTAGGCTGGAGCTGCTTC | GCGCAAGATCATGTTACCGGTAAAATAACCATAAAGGATAAGCGCAGATATTAACGGCTGACATGGGAAT | Integrated DNA Technologies, Inc. |  | | | | | |  |
| Δ*talA*::KAN(R) | CGCACTCATCTAACACTTTACTTTTCAAGGAGTATTTCCTGTGTAGGCTGGAGCTGCTTC | GGCAAGGTCTTTTCGGGACATATAACACTCCGTGGCTGGTTTAACGGCTGACATGGGAAT | Integrated DNA Technologies, Inc. |  | | | | | |  |
| **Oligonucleotides to Generate Plasmids Construction** | | | |  |  |  |  |  |  |  |
| **Plasmid/Deletion** | **Forward Primer (5’ to 3’)** | **Reverse Primer (5’ to 3’)** | **Source** |  | | | | | |  |
| pUA66-*crp* | GCGCTCAGATCTTGATCCGAAAGCTATGCTAAAACAGT | GCGCTCAGTACTttaACGAGTGCCGTAAACGA | Integrated DNA Technologies, Inc. |  | | | | | |  |
| **Oligonucleotides to Verify Gene Deletions** | | | |  |  |  |  |  |  |  |
| **Mutation** | **External Forward Primer (5’ to 3’)** | **External Reverse Primer (5’ to 3’)** | **Internal Forward Primer (5’ to 3’)** | **Internal Reverse Primer (5’ to 3’)** | | | | | | **Source** |
| Δ*crp*::KAN(R) | CAAAGCGAAAGCTATGCTAAAACAG | GCTCTTCGTCCAGATCATCCT | ACAGACCCGACTCTCGAATG | GAGTGCCGTAAACGACGATG | | | | | | Integrated DNA Technologies, Inc. |
| Δ*cyaA*::KAN(R) | CGCATCTTTCTTTACGGTCAATC | GCTCTTCGTCCAGATCATCCT | GCTAATGCCGGGTTACCTTG | CGCAAAGCGGATCAGATTAC | | | | | | Integrated DNA Technologies, Inc. |
| Δ*sucA*::KAN(R) | TGCCCCTGACACTAAGACAGT | GCTCTTCGTCCAGATCATCCT | TACCTCGGCGCAAAATTCCC | GACGACGCAGCATGTGGTAA | | | | | | Integrated DNA Technologies, Inc. |
| Δ*lpd*::KAN(R) | CCTATGGATTTCTGGGTGCAG | GCTCTTCGTCCAGATCATCCT | CTTCCGTTGCGCTGATTTAG | CAGCATCACAACCCATTTCG | | | | | | Integrated DNA Technologies, Inc. |
| Δ*sucC*::KAN(R) | CATTATCGCCTTCTCCGGCA | GCTCTTCGTCCAGATCATCCT | ACAACTTTTTGCCCGCTATG | GAGCTGCATCCGTCAGACCT | | | | | | Integrated DNA Technologies, Inc. |
| Δ*sdhA*::KAN(R) | CTGGAT CGGTTT CTTCGCCT | GCTCTTCGTCCAGATCATCCT | TTACCG TTGCGC TGGGTA AT | GTGAGC GAAGGT ACGGGA AA | | | | | | Integrated DNA Technologies, Inc. |
| Δ*gltA*::KAN(R) | AGGTTGATGTGCGAAGGCAA | GCTCTTCGTCCAGATCATCCT | ACCTTTGACCCAGGCTTCAC | GGAAGACGGAATACCCATCGC | | | | | | Integrated DNA Technologies, Inc. |
| Δ*acnB*::KAN(R) | TCCTGCTATTCTGCCCGTTG | GCTCTTCGTCCAGATCATCCT | GCGAAAGGCGAAGCCAAATC | GATCTCGATACGCGCACCAC | | | | | | Integrated DNA Technologies, Inc. |
| Δ*aceE*::KAN(R) | GCGCGGCAACTAAACGTAGA | GCTCTTCGTCCAGATCATCCT | TCCTATCCGCACCCGAAACT | GCTCGTTGATCCCTTCCTGC | | | | | | Integrated DNA Technologies, Inc. |
| Δ*fumA*::KAN(R) | GTAACCTGGAGCCGCAAAAA | GCTCTTCGTCCAGATCATCCT | CATTATCAGGCTCCTTTTCCAC | AAACGGGATACTGCGACAAC | | | | | | Integrated DNA Technologies, Inc. |
| Δ*mdh*::KAN(R) | CGCTAAACTTGCGTGACTACAC | GCTCTTCGTCCAGATCATCCT | CCCAACTGCCTTCAGGTTCAG | TGTTCAAATGCGCTCAGGGT | | | | | | Integrated DNA Technologies, Inc. |
| Δ*nuoI*::KAN(R) | GCCATTCATCTGGTTCGCGC | GCTCTTCGTCCAGATCATCCT | TTCGGCACCCAGGTTCGTAG | GGCTTCGTTCTCTGCTTCGC | | | | | | Integrated DNA Technologies, Inc. |
| Δ*nuoM*::KAN(R) | GCGTTGAGTTAAGGATTGTGG | GCTCTTCGTCCAGATCATCCT | CTTTATTGGCGGCTTCCTGT | CTGGCTGCCGGTGTAGATAG | | | | | | Integrated DNA Technologies, Inc. |
| Δ*atpA*::KAN(R) | GAAAATTTCTGCTGCGATGG | GCTCTTCGTCCAGATCATCCT | CACCGAAATCAGCGAACTG | GGATTGGGTTGCTTTGAAGG | | | | | | Integrated DNA Technologies, Inc. |
| Δ*atpB*::KAN(R) | TTGGTGTTACTGGTGGTGGC | GCTCTTCGTCCAGATCATCCT | GACGCCGCAGGATTACATAGG | TGCAGCGTCAACTCTTTCGT | | | | | | Integrated DNA Technologies, Inc. |
| Δ*atpC*::KAN(R) | GGCGAATACGATCACCTGCC | GCTCTTCGTCCAGATCATCCT | TAGCGAAGGTGAACTGGGGA | GTCAACTCGATAACGCGCAGC | | | | | | Integrated DNA Technologies, Inc. |
| Δ*atpD*::KAN(R) | CGGCAGCCTGATTAAAGAGC | GCTCTTCGTCCAGATCATCCT | CGTCGAATTCCCTCAGGATG | GCTTTTTCCACAGCTTCTTCG | | | | | | Integrated DNA Technologies, Inc. |
| Δ*frdC*::KAN(R) | TAAGAAGGAGCGTATGGCGC | GCTCTTCGTCCAGATCATCCT | GTTTTATCGCTTTTACATGCTGCG | AACAGGATTACGATGGTGGCAA | | | | | | Integrated DNA Technologies, Inc. |
| Δ*pgi*::KAN(R) | ACACTCAACATTACGCTAACGGC | GCTCTTCGTCCAGATCATCCT | GCGAGAAGATCAACCGCACT | TTCCTGCTCAACCACTTCGC | | | | | | Integrated DNA Technologies, Inc. |
| Δ*zwf*::KAN(R) | AATCGCACGGGTGGATAAGC | GCTCTTCGTCCAGATCATCCT | GGCGAGGCAAAACTGAATGC | TGATACGGTTTCGGCGCATC | | | | | | Integrated DNA Technologies, Inc. |
| Δ*talA*::KAN(R) | GAGCAAGTCCAAACTCTCACCATT | GCTCTTCGTCCAGATCATCCT | CCACCACCAATCCTTCGCTG | TTCTACCGCCATCGCATCCT | | | | | | Integrated DNA Technologies, Inc. |
| **Oligonucleotides to Verify Plasmids Construction** | | | |  |  |  |  |  |  |  |
| **Plasmid/Deletion** | **Forward Primer (5’ to 3’)** | **Reverse Primer (5’ to 3’)** | **Source** |  | | | | | |  |
| pUA66-*crp* | CGATCCTCATCCTGTCTCTT | CTGATCTTCCAGCATCTTCA | Integrated DNA Technologies, Inc. |  | | | | | |  |
| pUA66-EV | GCGCCTCATGAGGTACCCCGGGTCGACCTGCAGCCAAGCTTAATTA | GCGCCTCATGAACTAGAGGTCTCCTCTTTAATGAATTCTGTGTG | Integrated DNA Technologies, Inc. |  | | | | | |  |
