## Supplementary File 1 for "UNRAVELING CRP/cAMP-MEDIATED METABOLIC REGULATION IN *ESCHERICHIA COLI* PERSISTER CELLS"

**Supplementary File 1. MIC of antibiotics and concentrations of bactericidal antibiotics used in persister assays.**

|  | Concentration (μg/mL) | | |
| --- | --- | --- | --- |
| **Bacterial Strains** | **Ampicillin** | **Ofloxacin** | **Gentamicin** |
| MIC of *E. coli* K-12 MG1655 Wild Type | 3-4 | 0.032-0.047 | 0.19-0.25 |
| MIC of *E. coli* K-12 MG1655 Δ*crp* | 4-6 | 0.064-0.094 | 3-4 |
| MIC of *E. coli* K-12 MG1655 Δ*cyaA* | 6-8 | 0.064-0.094 | 3-4 |
| Persister Assay Concentration | 200 | 5 | 50 |
