## Supplementary File 2 for "UNRAVELING CRP/cAMP-MEDIATED METABOLIC REGULATION IN *ESCHERICHIA COLI* PERSISTER CELLS"

**Supplementary File 2. Analyzed metabolomics data.**

**Supplementary File 2A. Downregulated metabolites at early (t=5h) and late (t=24h) stationary phases with a threshold of 0.5 or lower for the ratio in the mutant *crp* to WT.**

| **Early stationary phase** |  |
| --- | --- |
| **Biochemical Name** | **Δ*crp*/WT** |
| 5,6-dihydrothymine | 0.01 |
| galactitol (dulcitol) | 0.01 |
| indole | 0.01 |
| urate | 0.01 |
| 2,3-dihydroxy-5-methylthio-4-pentenoate (DMTPA)* | 0.04 |
| 3-ureidopropionate | 0.04 |
| indolin-2-one | 0.04 |
| 2'-deoxycytidine 5'-monophosphate | 0.05 |
| N-acetylthreonine | 0.05 |
| pyridoxine phosphate | 0.05 |
| N-propionylmethionine | 0.06 |
| lysine | 0.07 |
| 3'-dephospho-CoA-glutathione* | 0.08 |
| gamma-glutamyl-2-aminobutyrate | 0.08 |
| chiro-inositol | 0.09 |
| gamma-glutamylalanine | 0.1 |
| palmitoleate (16:1n7) | 0.1 |
| deoxycarnitine | 0.11 |
| N-formylanthranilic acid | 0.11 |
| pseudouridine | 0.12 |
| pyridoxine (Vitamin B6) | 0.13 |
| 3'-dephospho-acetyl-CoA | 0.14 |
| threonate | 0.14 |
| 1,5-anhydroglucitol (1,5-AG) | 0.15 |
| 2'-deoxyinosine | 0.15 |
| cytidine 5'-monophosphate (5'-CMP) | 0.15 |
| mannitol-1-phosphate | 0.15 |
| N-succinyl-phenylalanine | 0.16 |
| 2-methylcitrate/homocitrate | 0.17 |
| 3'-dephosphocoenzyme A | 0.17 |
| cytosine | 0.17 |
| prolylglycine | 0.17 |
| riboflavin (Vitamin B2) | 0.18 |
| 2R,3R-dihydroxybutyrate | 0.19 |
| margarate (17:0) | 0.21 |
| kynurenate | 0.22 |
| myo-inositol | 0.22 |
| pyridoxate | 0.22 |
| 4-imidazoleacetate | 0.23 |
| phosphopantetheine | 0.23 |
| gamma-glutamylcysteine | 0.24 |
| phenylpyruvate | 0.24 |
| 4-hydroxyphenylacetate | 0.25 |
| 1-palmitoleoylglycerol (16:1)* | 0.26 |
| 3-methyl-2-oxobutyrate | 0.26 |
| 5-dodecenoate (12:1n7) | 0.27 |
| butyryl/isobutyryl CoA | 0.28 |
| 2-oxoarginine* | 0.3 |
| citrate | 0.3 |
| gamma-glutamylglycine | 0.3 |
| nicotinamide ribonucleotide (NMN) | 0.3 |
| nicotinic acid mononucleotide (NaMN) | 0.3 |
| 1-methyl-beta-carboline-3-carboxylic acid | 0.31 |
| 1-palmitoyl-GPE (16:0) | 0.31 |
| N-acetylhistidine | 0.31 |
| 1-palmitoyl-GPG (16:0)* | 0.32 |
| 4-hydroxyphenylpyruvate | 0.32 |
| arginine | 0.32 |
| argininosuccinate | 0.32 |
| 2'-deoxycytidine | 0.34 |
| 2-hydroxy-4-(methylthio)butanoic acid | 0.35 |
| glutamyl-meso-diaminopimelate | 0.35 |
| (3'-5')-adenylylguanosine* | 0.36 |
| (3'-5')-uridylylguanosine | 0.36 |
| 1-stearoyl-GPE (18:0) | 0.36 |
| 2-methylserine | 0.36 |
| homoserine | 0.37 |
| 2-hydroxybutyrate/2-hydroxyisobutyrate | 0.39 |
| 5-methylcytosine | 0.39 |
| arabonate/xylonate | 0.4 |
| methylsuccinate | 0.4 |
| pentadecanoate (15:0) | 0.4 |
| thymidine | 0.4 |
| 1-oleoyl-GPE (18:1) | 0.41 |
| 2-palmitoleoylglycerol (16:1)* | 0.42 |
| succinylglutamine | 0.42 |
| N6-(delta2-Isopentenyl)-adenine | 0.43 |
| N-succinyl-leucine | 0.43 |
| thiamin (Vitamin B1) | 0.43 |
| thymidine 5'-monophosphate | 0.44 |
| 4-methyl-2-oxopentanoate | 0.45 |
| pyridoxal phosphate | 0.45 |
| dihydroorotate | 0.46 |
| glutarate (C5-DC) | 0.46 |
| pyrraline | 0.46 |
| aconitate [cis or trans] | 0.47 |
| propionyl CoA | 0.47 |
| alanyl-glutamyl-meso-diaminopimelate | 0.48 |
| N-succinyl-isoleucine | 0.48 |
| oleate/vaccenate (18:1) | 0.5 |

| **Late stationary phase** |  |
| --- | --- |
| **Biochemical Name** | **Δ*crp*/WT** |
| indole | 0 |
| 4-imidazoleacetate | 0.01 |
| oleoyl ethanolamide | 0.01 |
| pantethine | 0.01 |
| 3-dehydroshikimate | 0.01 |
| 2,3-dihydroxy-5-methylthio-4-pentenoate (DMTPA)* | 0.02 |
| gamma-glutamylcitrulline* | 0.02 |
| sedoheptulose-7-phosphate | 0.02 |
| ribose | 0.02 |
| mannitol-1-phosphate | 0.02 |
| palmitoyl ethanolamide | 0.02 |
| thiamin (Vitamin B1) | 0.02 |
| N-formylanthranilic acid | 0.03 |
| cyclic dGSH (1) | 0.03 |
| 1-methylguanine | 0.03 |
| indolin-2-one | 0.03 |
| riboflavin (Vitamin B2) | 0.04 |
| 2,3-dihydroxyisovalerate | 0.04 |
| azetidine-2-carboxylic acid | 0.04 |
| arabonate/xylonate | 0.05 |
| sedoheptulose | 0.05 |
| malate | 0.05 |
| phosphopantetheine | 0.05 |
| enterolactone sulfate | 0.05 |
| 4-vinylguaiacol glucuronide | 0.05 |
| phenylpyruvate | 0.06 |
| 4-hydroxyphenylacetate | 0.06 |
| 4-hydroxyphenylpyruvate | 0.06 |
| N-palmitoylglycine | 0.06 |
| 3-ureidopropionate | 0.06 |
| pantoate | 0.06 |
| 2-hydroxy-3-methylvalerate | 0.07 |
| ribonate | 0.07 |
| arabitol/xylitol | 0.07 |
| 10-undecenoate (11:1n1) | 0.07 |
| 3-methyl-2-oxovalerate | 0.08 |
| ribitol | 0.08 |
| pantetheine | 0.08 |
| 3-deoxyoctulosonate | 0.08 |
| gentisate | 0.09 |
| 4-methyl-2-oxopentanoate | 0.09 |
| 2R,3R-dihydroxybutyrate | 0.09 |
| thiamin monophosphate | 0.09 |
| carboxyethyl-GABA | 0.1 |
| 3-(4-hydroxyphenyl)lactate | 0.1 |
| 2-oxoarginine* | 0.1 |
| argininate* | 0.1 |
| UDP-N-acetylmuraminate (UDP-MurNAc)* | 0.1 |
| 3'-dephospho-CoA-glutathione* | 0.11 |
| 7-methylguanine | 0.11 |
| 4-thiouracil | 0.11 |
| phenylacetate | 0.12 |
| anthranilate | 0.12 |
| 3-methyl-2-oxobutyrate | 0.12 |
| N1,N12-diacetylspermine | 0.12 |
| pyridoxate | 0.12 |
| N-propionylmethionine | 0.13 |
| gamma-glutamyl-2-aminobutyrate | 0.13 |
| 5-(2-Hydroxyethyl)-4-methylthiazole | 0.13 |
| glutamyl-meso-diaminopimelate | 0.13 |
| indolelactate | 0.14 |
| citrate | 0.14 |
| 1-stearoyl-GPG (18:0) | 0.15 |
| 1-heptadecenoylglycerol (17:1)* | 0.15 |
| prolylglycine | 0.17 |
| ethyl alpha-glucopyranoside | 0.17 |
| pyroglutamine* | 0.18 |
| phenyllactate (PLA) | 0.18 |
| N-acetylglucosamine/N-acetylgalactosamine | 0.18 |
| galactitol (dulcitol) | 0.19 |
| erythronate* | 0.19 |
| aconitate [cis or trans] | 0.19 |
| 4-ethylphenylsulfate | 0.19 |
| gluconate | 0.19 |
| argininosuccinate | 0.2 |
| 1,5-anhydroglucitol (1,5-AG) | 0.2 |
| 3-formylindole | 0.2 |
| 3-indoleglyoxylic acid | 0.2 |
| pseudouridine | 0.21 |
| gulonate* | 0.21 |
| pyridoxine phosphate | 0.22 |
| N,N,N-trimethyl-5-aminovalerate | 0.23 |
| urate | 0.23 |
| S-carboxymethyl-L-cysteine | 0.23 |
| alpha-hydroxyisocaproate | 0.24 |
| alpha-hydroxyisovalerate | 0.24 |
| 3'-dephosphocoenzyme A | 0.24 |
| serine | 0.25 |
| 1-oleoyl-GPG (18:1)* | 0.25 |
| pipecolate | 0.26 |
| 5-methylcytosine | 0.26 |
| pyridoxal | 0.26 |
| fumarate | 0.27 |
| 2-hydroxyadipate | 0.27 |
| 2'-O-methylguanosine | 0.27 |
| adenine | 0.28 |
| N6-methyllysine | 0.29 |
| kynurenate | 0.29 |
| 3-indoxyl sulfate | 0.29 |
| (12 or 13)-methylmyristate (a15:0 or i15:0) | 0.29 |
| glutarate (C5-DC) | 0.29 |
| hypoxanthine | 0.29 |
| nicotinate | 0.29 |
| indole-3-carboxylate | 0.3 |
| 2-hydroxy-4-(methylthio)butanoic acid | 0.3 |
| 3-ethylphenylsulfate | 0.3 |
| 4-hydroxy-2-oxoglutaric acid | 0.31 |
| 1-stearoyl-GPE (18:0) | 0.33 |
| N6-dimethylallyladenine | 0.33 |
| sucrose | 0.34 |
| N-acetylglutamate | 0.35 |
| 2-hydroxybutyrate/2-hydroxyisobutyrate | 0.35 |
| maleate | 0.35 |
| 1-palmitoyl-2-oleoyl-GPE (16:0/18:1) | 0.35 |
| choline phosphate | 0.36 |
| pyrraline | 0.37 |
| 2-aminoadipate | 0.38 |
| 4-acetamidobutanoate | 0.38 |
| phosphoethanolamine | 0.38 |
| 3'-dephospho-acetyl-CoA | 0.38 |
| salicylate | 0.38 |
| glycylleucine | 0.39 |
| ribulonate/xylulonate/lyxonate* | 0.39 |
| 1-stearoyl-2-linoleoyl-GPE (18:0/18:2)* | 0.39 |
| prephenic acid | 0.4 |
| pyridoxal phosphate | 0.4 |
| 4-methylcatechol sulfate | 0.4 |
| beta-guanidinopropanoate | 0.4 |
| acetylphosphate | 0.41 |
| myo-inositol | 0.41 |
| 1-oleoyl-GPE (18:1) | 0.41 |
| cytosine | 0.41 |
| mannonate* | 0.41 |
| flavin mononucleotide (FMN) | 0.42 |
| 4-ethylcatechol sulfate | 0.42 |
| equol sulfate | 0.42 |
| 1-carboxyethylphenylalanine | 0.43 |
| N-succinyl-isoleucine | 0.43 |
| 4-guanidinobutanoate | 0.43 |
| o-Tyrosine | 0.44 |
| methylsuccinate | 0.44 |
| 5-oxoproline | 0.44 |
| 1,2-dioleoyl-GPE (18:1/18:1) | 0.44 |
| diadenosine triphosphate | 0.44 |
| 2-methylcitrate/homocitrate | 0.45 |
| margarate (17:0) | 0.45 |
| benzoate | 0.46 |
| isobutyrylglycine | 0.47 |
| homocitrulline | 0.47 |
| 1-stearoyl-2-oleoyl-GPE (18:0/18:1) | 0.47 |
| imidazole lactate | 0.48 |
| (14 or 15)-methylpalmitate (a17:0 or i17:0) | 0.48 |
| 2'-O-methyluridine | 0.48 |
| pyridoxine (Vitamin B6) | 0.48 |
| valylglycine | 0.49 |
| glucose | 0.49 |
| erucate (22:1n9) | 0.49 |
| N-carbamoylvaline | 0.5 |
| 1-palmitoleoylglycerol (16:1)* | 0.5 |
| thymine | 0.5 |

**Supplementary File 2B. Upregulated metabolites at early (t=5h) and late (t=24h) stationary phases with a threshold of 2 or higher for the ratio in the mutant *crp* to WT.**

| **Early stationary phase** |  |
| --- | --- |
| **Biochemical Name** | **Δ*crp*/WT** |
| thioproline | 169.74 |
| N-acetylcysteine | 168.31 |
| cysteine | 157.41 |
| N-acetylneuraminate | 141.67 |
| Fructose 1,6-diphosphate/glucose 1,6-diphosphate/myo-inositol diphosphates | 126.81 |
| maltotriose | 89.56 |
| cysteine-glutathione disulfide | 76.98 |
| xanthosine | 74.39 |
| gamma-glutamylleucine | 70.22 |
| maltose | 54.06 |
| maltotetraose | 48.43 |
| mannonate* | 38 |
| 3-phosphoglycerate | 26.38 |
| dihydroxyacetone phosphate (DHAP) | 24.77 |
| cysteine sulfinic acid | 23.54 |
| methylmalonate (MMA) | 22.92 |
| gamma-glutamyltyrosine | 21.96 |
| acetylagmatine | 20.92 |
| 2-phosphoglycerate | 20.59 |
| gamma-glutamylphenylalanine | 19.74 |
| erythritol | 17.44 |
| agmatine | 16.99 |
| glycerate | 16.38 |
| asparagine | 15.21 |
| acetylphosphate | 14.82 |
| gamma-glutamylmethionine | 14.21 |
| glucuronate | 14.05 |
| ergothioneine | 13.23 |
| 3-phosphoserine | 12.57 |
| UDP-glucuronate | 11.27 |
| gamma-glutamylhistidine | 10 |
| 2'-deoxyadenosine 5'-diphosphate | 9.34 |
| 5-methyluridine (ribothymidine) | 9.31 |
| uracil | 8.65 |
| 3-sulfo-L-alanine | 8.44 |
| sorbitol 6-phosphate | 8.33 |
| UDP-galactose | 7.95 |
| phosphoenolpyruvate (PEP) | 7.72 |
| lysylleucine | 7.67 |
| UDP-glucose | 7.6 |
| 4-acetamidobutanoate | 7.59 |
| gamma-glutamylglutamate | 6.66 |
| histamine | 6.61 |
| uridine 5'-diphosphate (UDP) | 6.61 |
| 5-aminovalerate | 6.54 |
| gluconate | 6.46 |
| threonine | 6.27 |
| fructose | 6.11 |
| lactobacillic acid | 6.02 |
| valylleucine | 5.86 |
| tryptophan | 5.84 |
| N-acetylglucosamine 6-phosphate | 5.82 |
| gamma-glutamylisoleucine* | 5.77 |
| mannitol/sorbitol | 5.7 |
| 4-hydroxybutyrate (GHB) | 5.67 |
| beta-alanine | 5.67 |
| lactate | 5.49 |
| aspartate | 5.46 |
| 1,3-diaminopropane | 5.42 |
| N-acetyl-cadaverine | 5.3 |
| orotate | 5.26 |
| 2-heptadecenoylglycerol (17:1)* | 4.76 |
| 2'-O-methyluridine | 4.67 |
| ribulose/xylulose | 4.67 |
| 3-hydroxyadipate* | 4.61 |
| guanosine-2',3'-cyclic monophosphate | 4.51 |
| N6,N6-dimethyladenosine | 4.42 |
| thiamin diphosphate | 4.39 |
| diaminopimelate | 4.37 |
| indolelactate | 4.25 |
| uridine | 4.15 |
| N-acetylglutamate | 3.81 |
| 1,2-dipalmitoyl-GPC (16:0/16:0) | 3.8 |
| diacetylspermidine* | 3.78 |
| serine | 3.75 |
| 2-Dehydro-3-deoxy-D-gluconate | 3.68 |
| alanylleucine | 3.67 |
| gamma-glutamyltryptophan | 3.63 |
| N-acetylaspartate (NAA) | 3.56 |
| nicotinate ribonucleoside | 3.53 |
| adenosine 3',5'-cyclic monophosphate (cAMP) | 3.49 |
| cadaverine | 3.44 |
| glycine | 3.43 |
| 5,6-dihydrouridine | 3.42 |
| glycylvaline | 3.39 |
| adenine | 3.24 |
| leucylglycine | 3.22 |
| Ectoine | 3.22 |
| spermine | 3.21 |
| cysteinylglycine | 3.06 |
| malate | 2.95 |
| homocysteine | 2.82 |
| ribitol | 2.8 |
| 2'-deoxyuridine | 2.79 |
| 3-hydroxymyristate | 2.67 |
| 1-heptadecenoylglycerol (17:1)* | 2.66 |
| heptanoate (7:0) | 2.63 |
| N1,N12-diacetylspermine | 2.58 |
| nicotinamide riboside | 2.55 |
| sebacate (C10-DC) | 2.49 |
| 3-hydroxybutyrate (BHBA) | 2.49 |
| 1-carboxyethylvaline | 2.45 |
| glutamate | 2.39 |
| guanosine 5'- monophosphate (5'-GMP) | 2.35 |
| glutathione, reduced (GSH) | 2.33 |
| (2 or 3)-decenoate (10:1n7 or n8) | 2.32 |
| inosine | 2.32 |
| guanosine | 2.31 |
| 4-hydroxybenzoate | 2.27 |
| N('1)-acetylspermidine | 2.26 |
| arachidate (20:0) | 2.2 |
| nicotinate | 2.18 |
| glutamate, gamma-methyl ester | 2.12 |
| glycerol 3-phosphate | 2.06 |
| erucate (22:1n9) | 2.02 |
| stachydrine | 2.01 |

| **Late stationary phase** |  |
| --- | --- |
| **Biochemical Name** | **Δ*crp*/WT** |
| N-acetylneuraminate | 106.91 |
| gamma-glutamyltryptophan | 72.05 |
| cysteine | 60.93 |
| xanthosine | 48.65 |
| ectoine | 40.72 |
| N2-acetyllysine | 39.34 |
| cysteine-glutathione disulfide | 36.07 |
| N-acetylcysteine | 33.59 |
| thioproline | 29.8 |
| maltotriose | 22.08 |
| agmatine | 18.61 |
| gamma-glutamylleucine | 17.16 |
| adenosine 3',5'-cyclic monophosphate (cAMP) | 15.58 |
| gamma-glutamylmethionine | 14.78 |
| thymidine 5'-monophosphate | 12.06 |
| glutamate, gamma-methyl ester | 11.46 |
| N-acetylmethionine | 11.23 |
| maltose | 10.29 |
| maltotetraose | 10.18 |
| gamma-glutamylhistidine | 9.09 |
| 3-hydroxypalmitate | 8.98 |
| 2'-deoxyguanosine 5'-monophosphate (dGMP) | 8.82 |
| diaminopimelate | 8.4 |
| beta-alanine | 8.36 |
| acetylagmatine | 8.31 |
| gamma-glutamylglutamine | 8.25 |
| neopterin | 8.23 |
| N-acetylisoleucine | 8.08 |
| gamma-glutamyltyrosine | 7.88 |
| N-acetylarginine | 7.15 |
| gamma-glutamylisoleucine* | 6.72 |
| glutathione, reduced (GSH) | 6.29 |
| glutamate | 5.95 |
| alanyl-glutamyl-meso-diaminopimelate | 5.86 |
| pyruvate | 5.83 |
| glycerol 3-phosphate | 5.81 |
| gamma-glutamylthreonine | 5.75 |
| 2'-deoxycytidine 5'-monophosphate | 5.61 |
| gamma-glutamylphenylalanine | 5.6 |
| 3-methylhistidine | 5.59 |
| cysteinylglycine | 5.59 |
| N-acetylvaline | 5.41 |
| homocysteine | 5.34 |
| 2'-deoxyadenosine 5'-monophosphate | 5.29 |
| deoxythymidine diphosphate-l-rhamnose | 5.15 |
| gamma-glutamylglutamate | 5.13 |
| 3-hydroxymyristate | 5.1 |
| ornithine | 5.06 |
| guanosine | 4.64 |
| gamma-glutamyl-epsilon-lysine | 4.61 |
| tryptophan | 4.35 |
| stachydrine | 4.35 |
| 2'-deoxyadenosine 5'-diphosphate | 4.31 |
| myristate (14:0) | 4.21 |
| uridine 5'-monophosphate (UMP) | 4.19 |
| acetyl CoA | 4.11 |
| alpha-ketoglutarate | 4.04 |
| gamma-glutamylglycine | 4 |
| glycerophosphoethanolamine | 3.99 |
| kynurenine | 3.94 |
| urea | 3.92 |
| UDP-galactose | 3.92 |
| UDP-glucose | 3.89 |
| cyclic(AMP-GMP) | 3.87 |
| guanosine-2',3'-cyclic monophosphate | 3.86 |
| 3-hydroxyoleate* | 3.78 |
| nicotinamide adenine dinucleotide (NAD+) | 3.66 |
| 3-hydroxyoctanoate | 3.65 |
| lysine | 3.62 |
| asparagine | 3.61 |
| N6-acetyllysine | 3.54 |
| 3-hydroxybutyrate (BHBA) | 3.54 |
| pantothenate | 3.43 |
| 2-methylserine | 3.37 |
| phenethylamine | 3.31 |
| erythritol | 3.31 |
| trimethylamine N-oxide | 3.23 |
| coenzyme A | 3.23 |
| N-acetylhistidine | 3.18 |
| gamma-glutamylserine | 3.11 |
| guanosine 5'- monophosphate (5'-GMP) | 3.1 |
| 9,10-methylenehexadecanoate** | 3.1 |
| R-methylmalonyl CoA | 3.07 |
| 2,4-di-tert-butylphenol | 3.06 |
| 3-hydroxylaurate | 3.03 |
| glucuronate | 2.98 |
| 1-palmitoleoyl-2-oleoyl-GPG (16:1/18:1)* | 2.89 |
| carnitine | 2.87 |
| 3-hydroxydecanoate | 2.86 |
| glycylvaline | 2.84 |
| uridine | 2.78 |
| cytidine | 2.78 |
| nicotinamide ribonucleotide (NMN) | 2.78 |
| palmitoleate (16:1n7) | 2.77 |
| N-formylmethionine | 2.76 |
| N-acetylmethionine sulfoxide | 2.76 |
| valylglutamine | 2.73 |
| N-acetylleucine | 2.72 |
| 3-amino-2-piperidone | 2.72 |
| methylmalonate (MMA) | 2.71 |
| N-acetylglucosamine 6-phosphate | 2.69 |
| adenosine 5'-diphosphoribose (ADP-ribose) | 2.67 |
| putrescine | 2.65 |
| 3-hydroxyhexanoate | 2.65 |
| pterin | 2.62 |
| fructose | 2.56 |
| N-acetylaspartate (NAA) | 2.5 |
| 4-hydroxybutyrate (GHB) | 2.48 |
| cysteine sulfinic acid | 2.46 |
| homoserine | 2.45 |
| N6,N6,N6-trimethyllysine | 2.41 |
| histamine | 2.39 |
| lactobacillic acid | 2.37 |
| lysylleucine | 2.33 |
| sebacate (C10-DC) | 2.33 |
| glycine | 2.28 |
| N-acetylputrescine | 2.28 |
| (2 or 3)-decenoate (10:1n7 or n8) | 2.27 |
| dimethylarginine (SDMA + ADMA) | 2.25 |
| UDP-glucuronate | 2.25 |
| orotate | 2.24 |
| UDP-N-acetylglucosamine/galactosamine | 2.21 |
| methionine | 2.15 |
| oleate/vaccenate (18:1) | 2.15 |
| succinyl CoA | 2.14 |
| 3-hydroxystearate | 2.14 |
| glutathione, oxidized (GSSG) | 2.06 |
