## Supplementary File 3 for "UNRAVELING CRP/cAMP-MEDIATED METABOLIC REGULATION IN *ESCHERICHIA COLI* PERSISTER CELLS"

**Supplementary File 3. Analyzed proteomics data.**

**Supplementary File 3A. The pathway analysis for the upregulated proteins in the Δ*crp* strain compared to WT.** This analysis integrates statistical analysis across the entire genome and includes various functional pathway classification frameworks such as Gene Ontology annotations, KEGG pathways, and Uniprot keywords. Count in Network: The first number indicates how many proteins in our network are annotated with a particular term. The second number indicates how many proteins in total (in our network and in the background) have this term assigned. Strength: Log_10_(observed/expected). This measure describes how large the enrichment effect is. It’s the ratio between i) the number of proteins in our network that are annotated with a term and ii) the number of proteins that we expect to be annotated with this term in a random network of the same size. False Discovery Rate: This measure describes how significant the enrichment is. Shown are p-values corrected for multiple testing within each category using the Benjamini–Hochberg procedure.

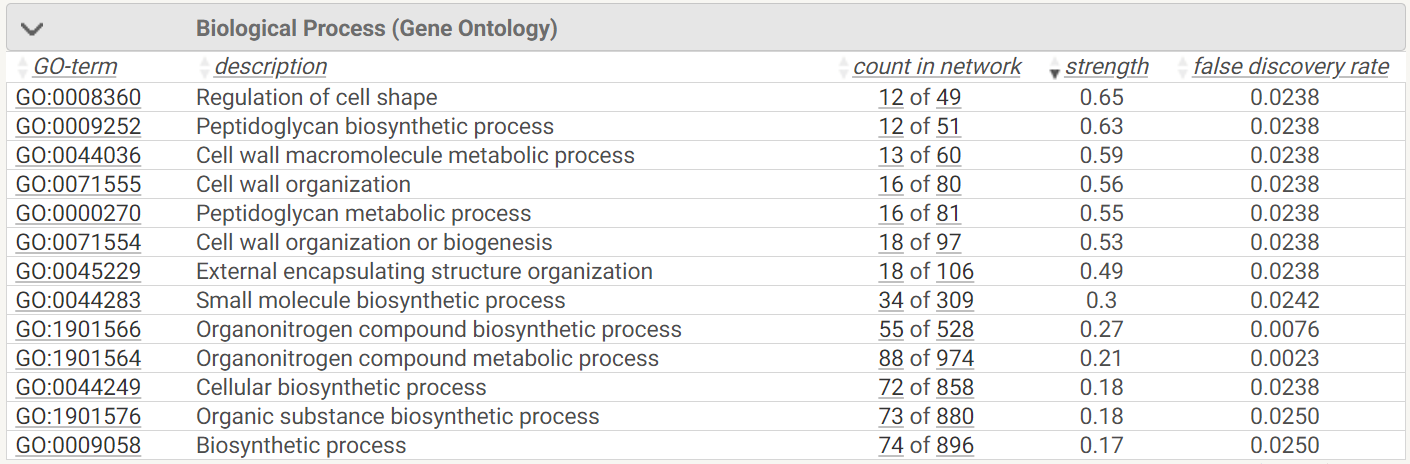

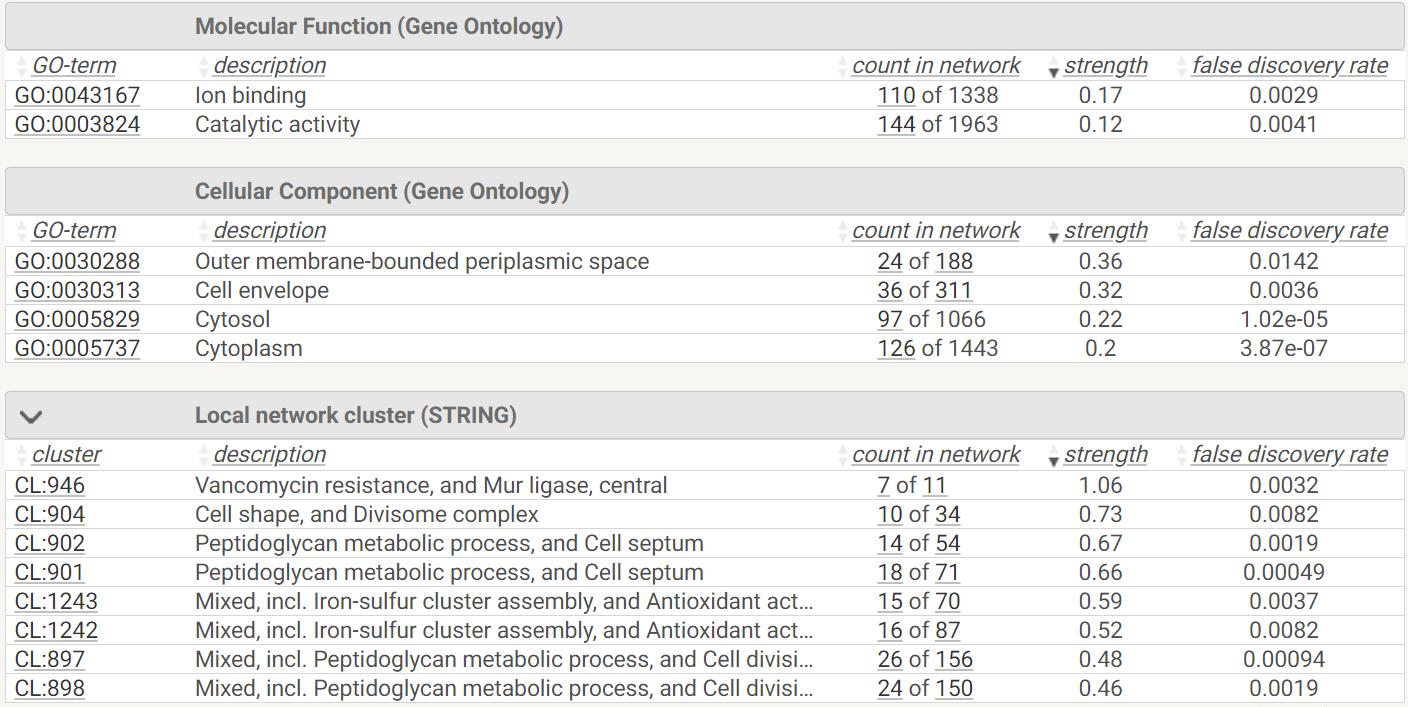

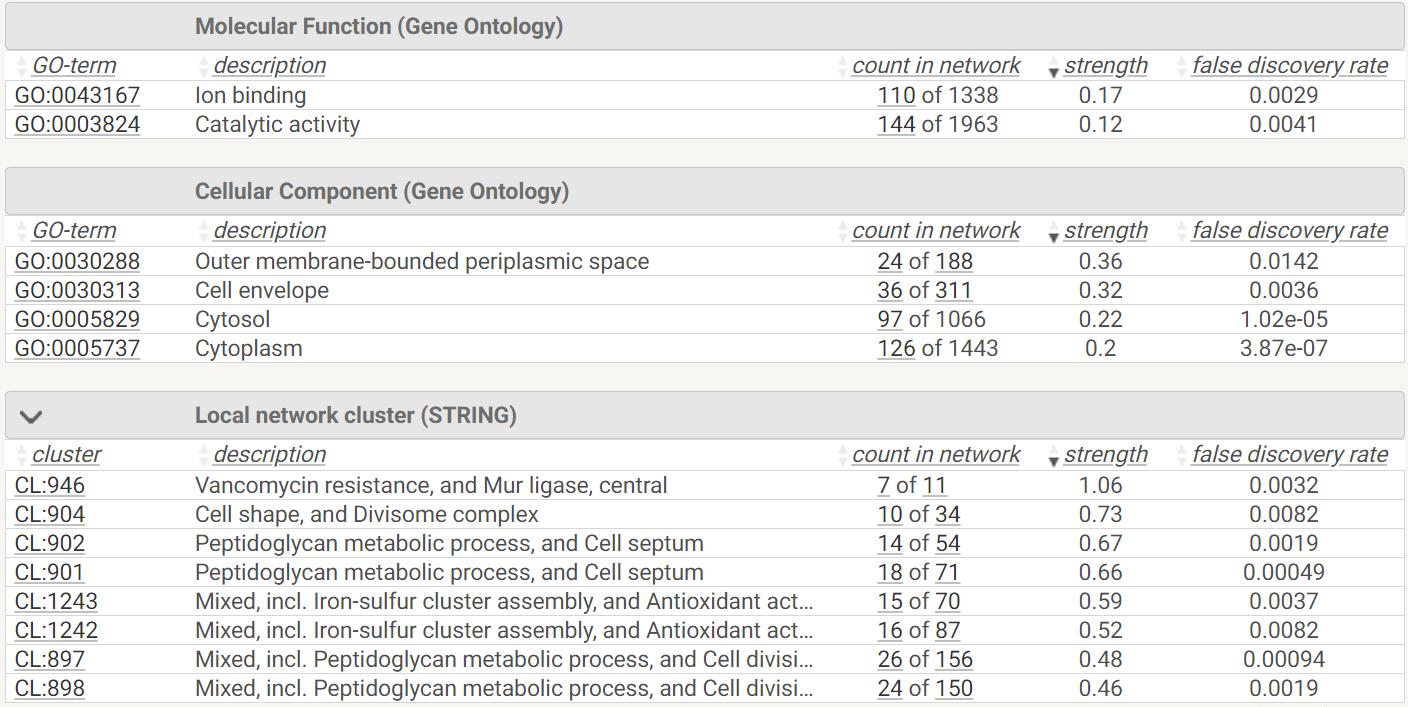

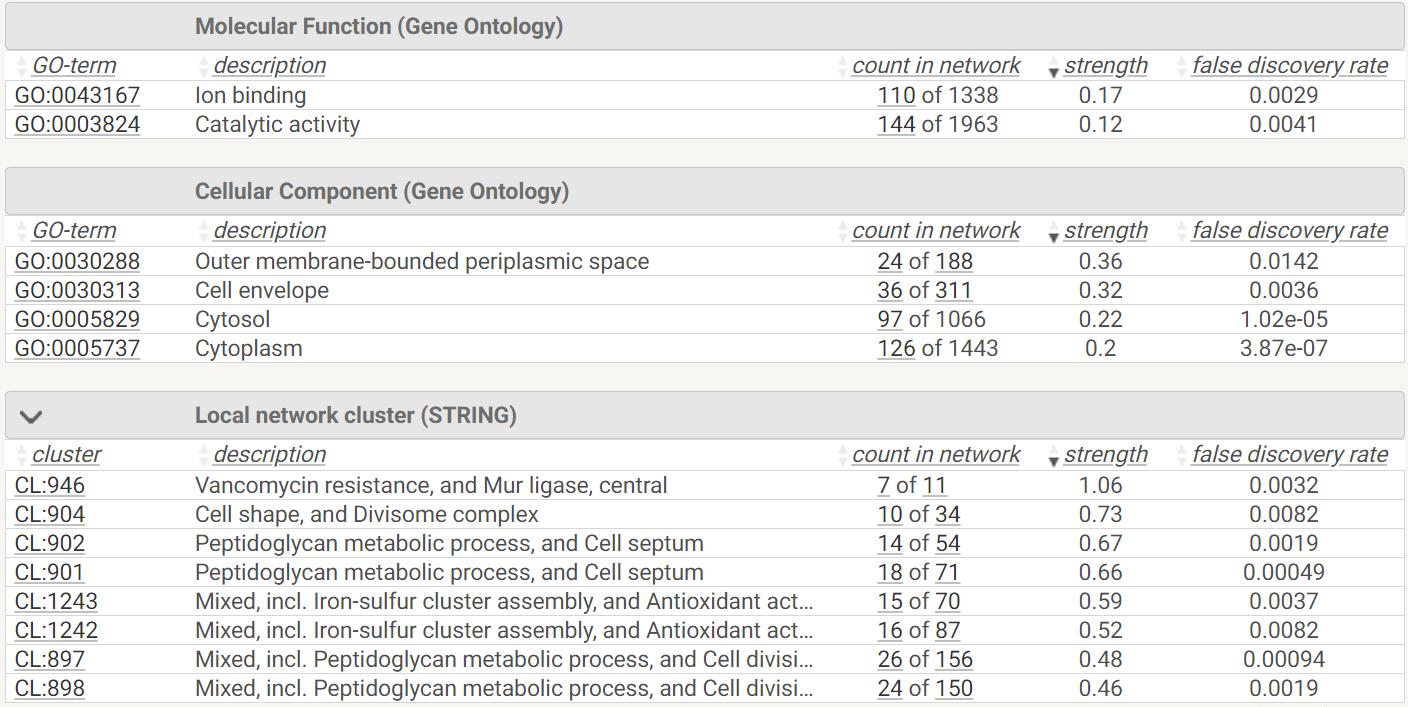

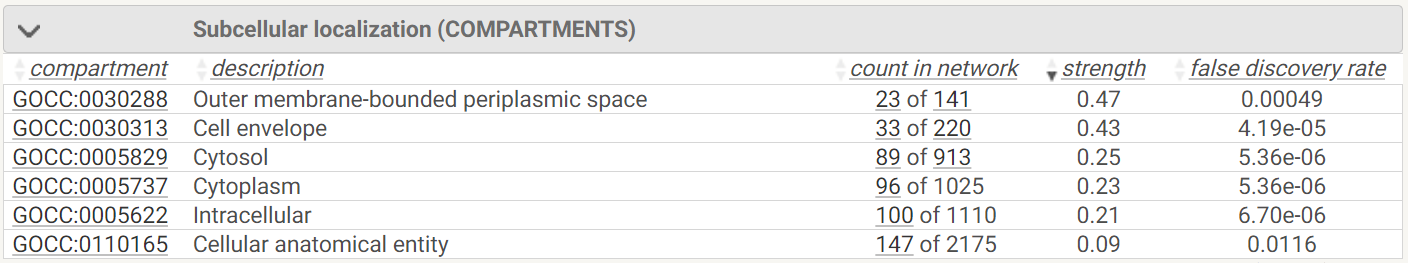

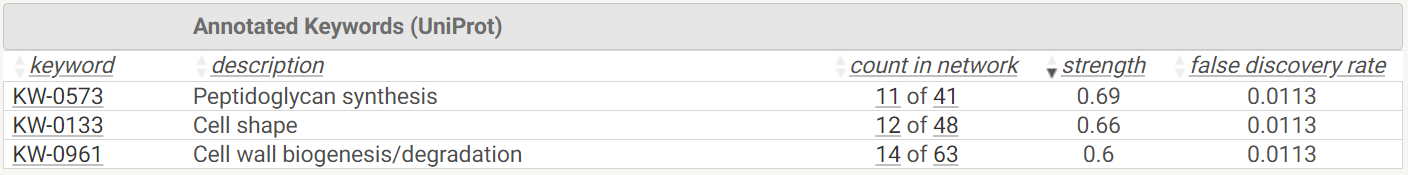

**Supplementary File 3B.** **The pathway analysis for the downregulated proteins in the Δ*crp* strain compared to WT.** See Supplementary File 3A for details.

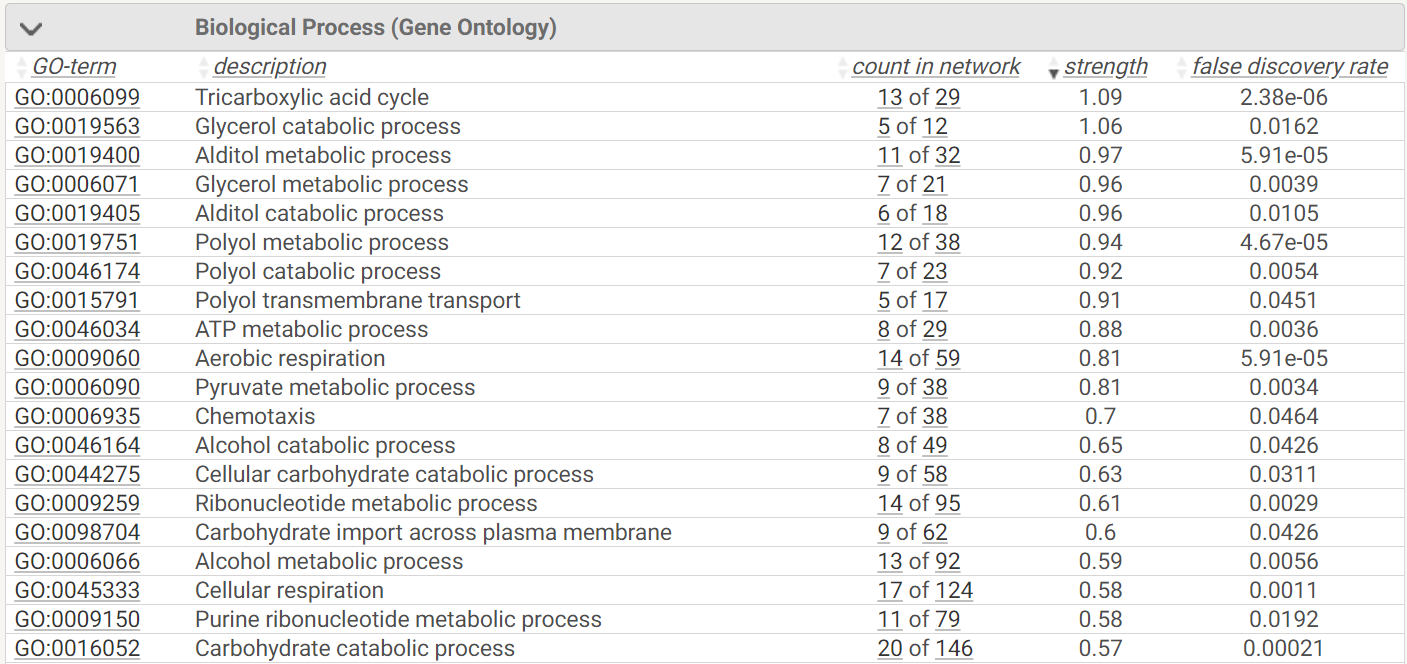

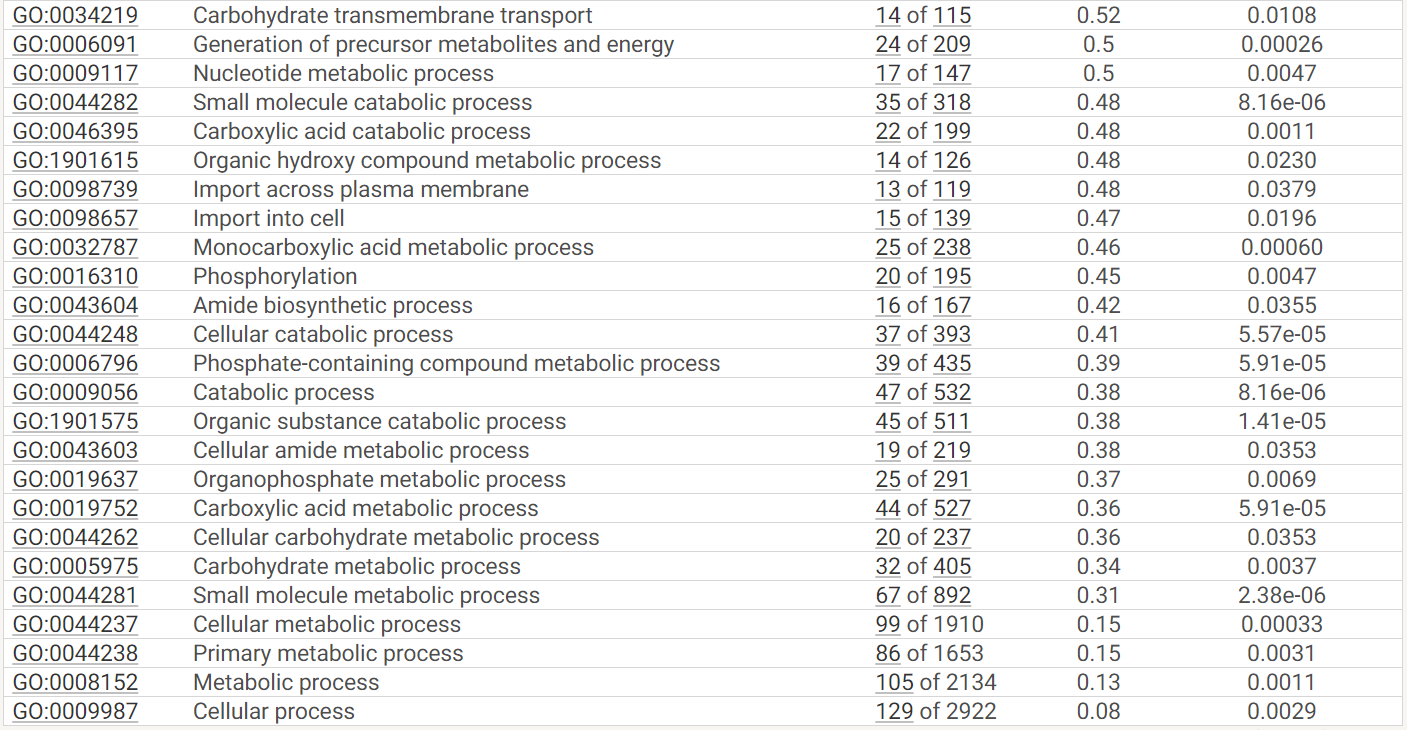

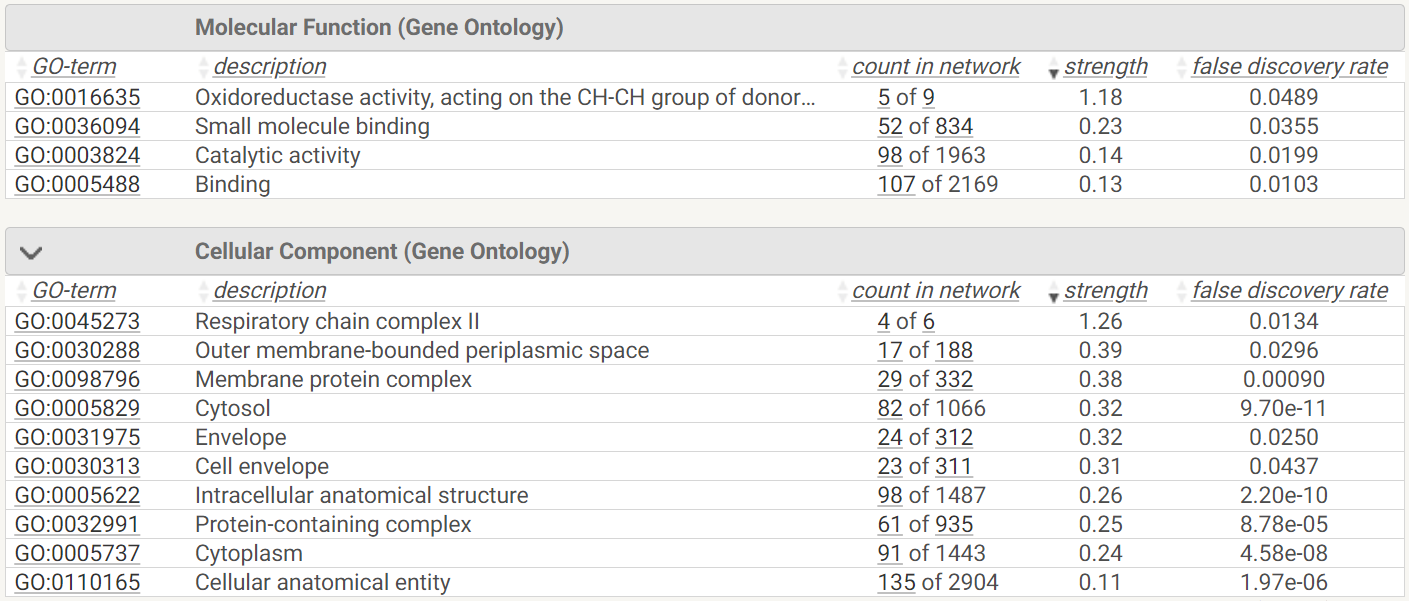

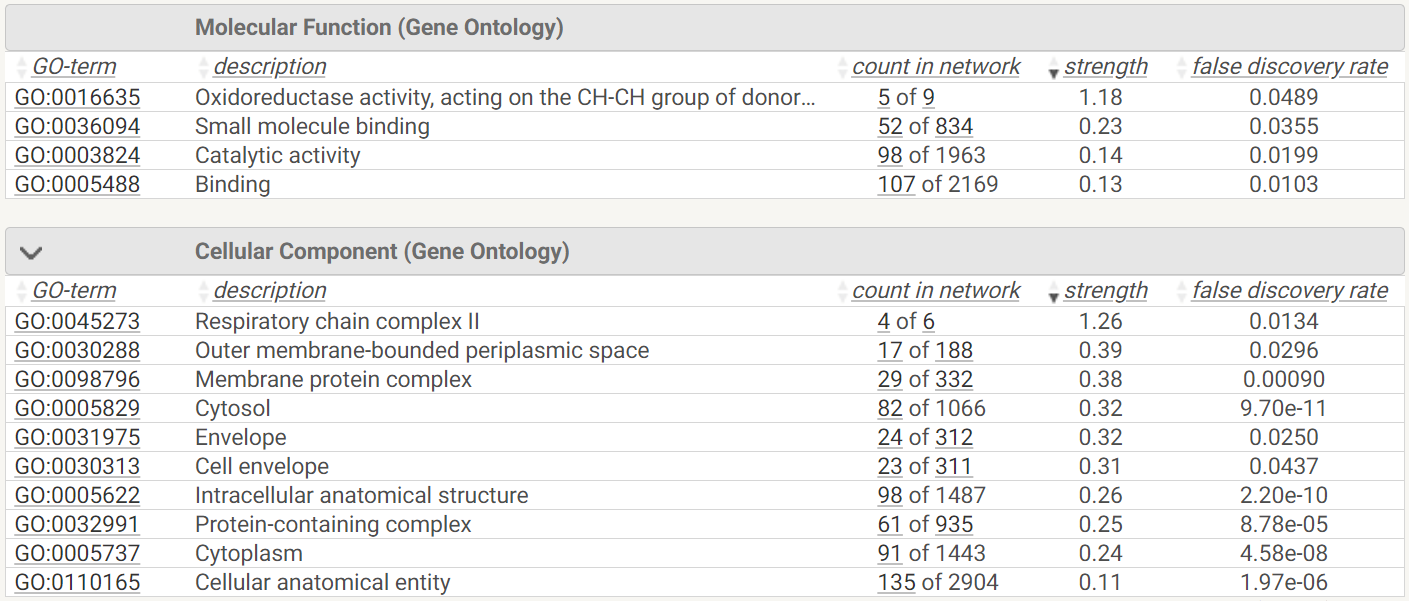

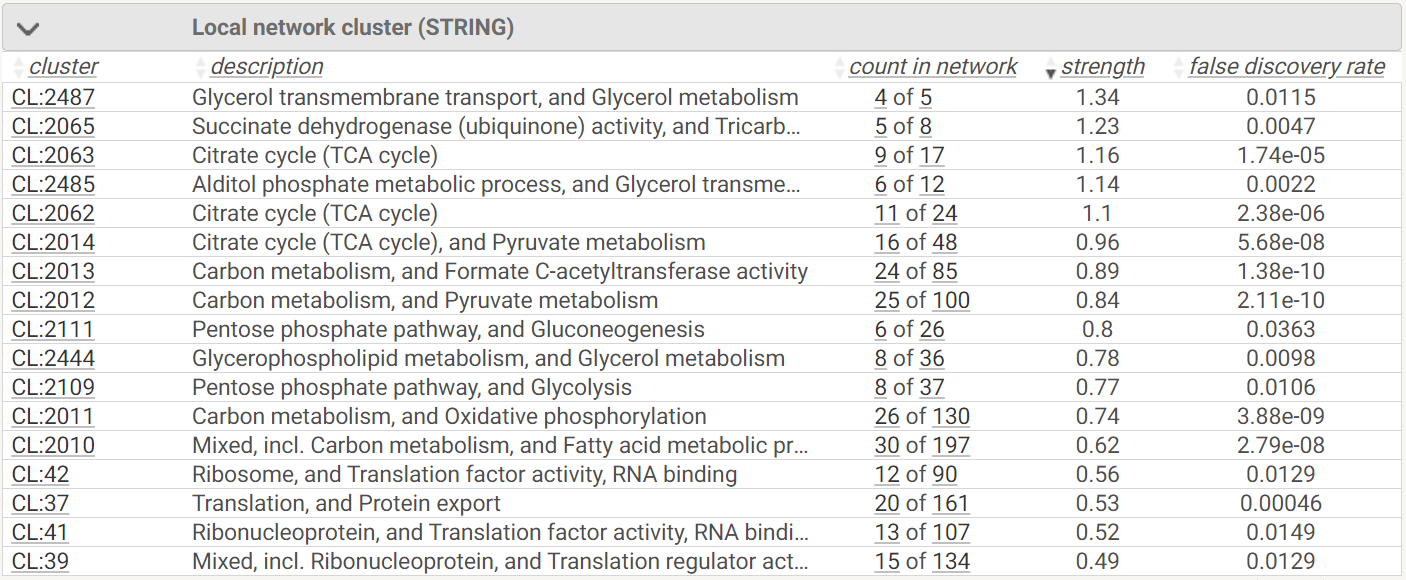

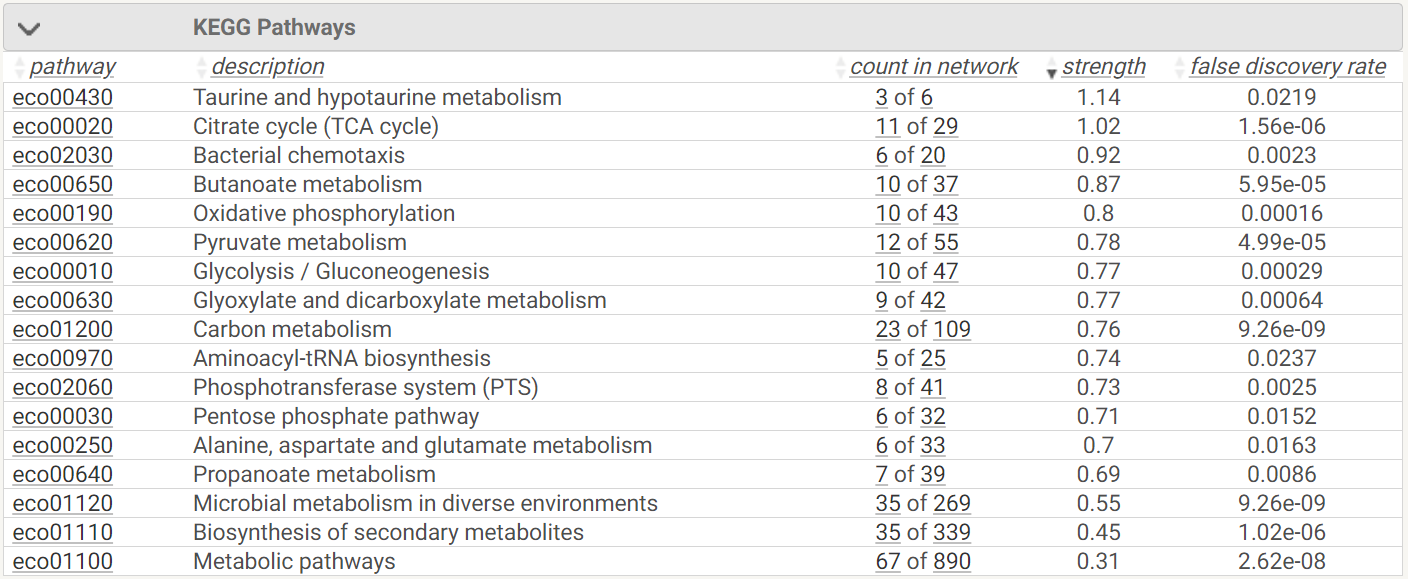

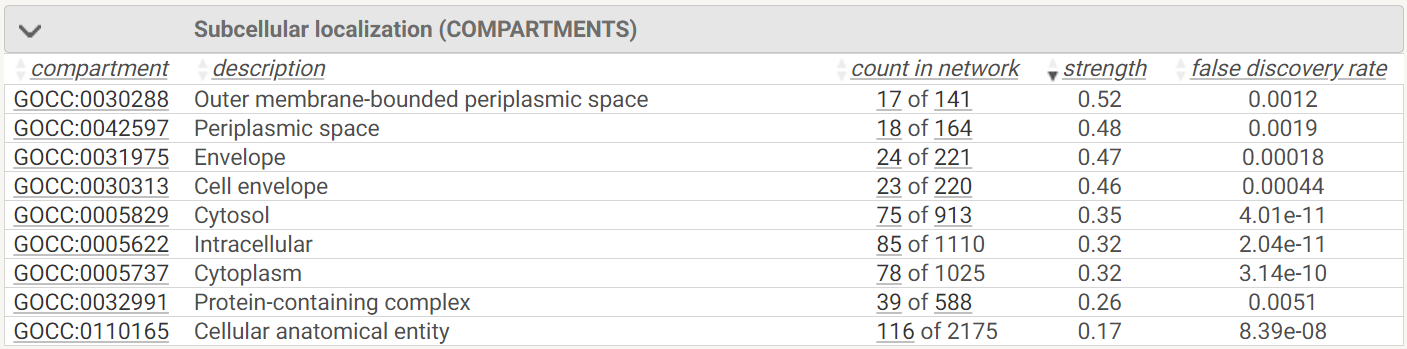

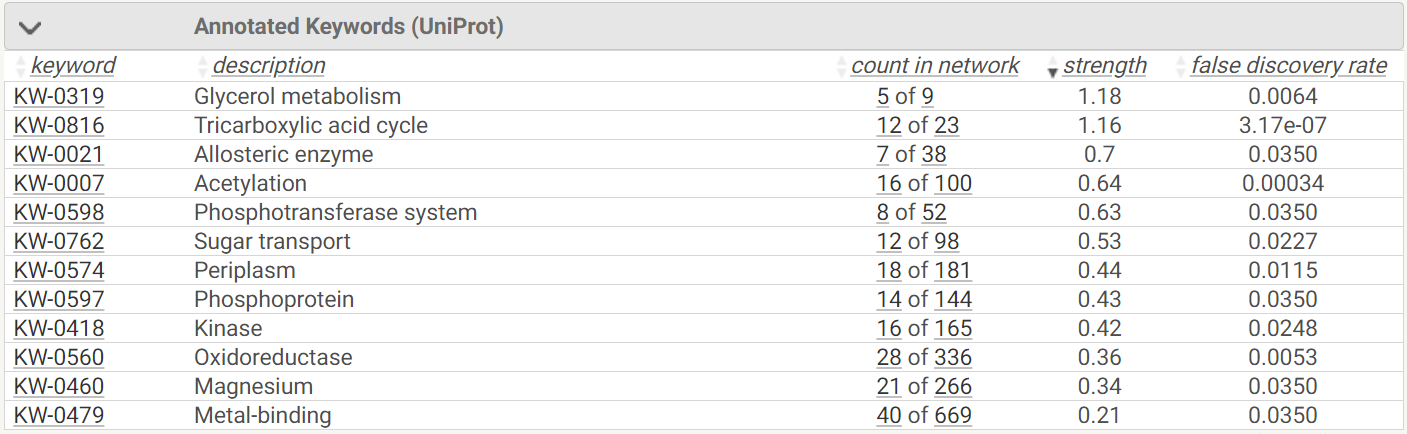
