## Supplementary File 4 for "UNRAVELING CRP/cAMP-MEDIATED METABOLIC REGULATION IN *ESCHERICHIA COLI* PERSISTER CELLS"

**Supplementary File 4. Analyzed persister survival fraction data.**

**Ampicillin and ofloxacin persister fractions.** Strains were from the single knockout (KO) Keio collection.

| KO strain | Ampicillin survival fraction |  | KO strain | Ofloxacin survival fraction |
| --- | --- | --- | --- | --- |
| *ΔldhA* | 2.94E-08 |  | *ΔgpmM* | 5.26E-08 |
| *ΔatpF* | 3.85E-08 |  | *ΔnuoM* | 5.88E-08 |
| *ΔnuoC* | 3.85E-08 |  | *ΔnuoI* | 7.69E-08 |
| *ΔfdhF* | 4.00E-08 |  | *ΔpfkA* | 8.00E-08 |
| *ΔnarV* | 4.35E-08 |  | *ΔaceE* | 8.70E-08 |
| *ΔnuoI* | 5.88E-08 |  | *ΔatpF* | 1.43E-07 |
| *ΔnuoM* | 6.67E-08 |  | *ΔappC* | 1.85E-07 |
| *ΔmaeB* | 6.90E-08 |  | *ΔnarV* | 2.78E-07 |
| *ΔsucB* | 9.09E-08 |  | *ΔfumA* | 3.33E-07 |
| *ΔgpmM* | 1.05E-07 |  | *ΔfdhF* | 3.33E-07 |
| *ΔaceE* | 1.18E-07 |  | *ΔatpC* | 5.26E-07 |
| *ΔsucD* | 1.33E-07 |  | *ΔsdhA* | 5.77E-07 |
| *ΔsdhC* | 1.64E-07 |  | *ΔfumC* | 6.00E-07 |
| *ΔnuoB* | 2.11E-07 |  | *ΔmaeB* | 6.92E-07 |
| *ΔsucA* | 2.17E-07 |  | *ΔsdhC* | 1.00E-06 |
| *ΔsdhA* | 2.17E-07 |  | *ΔsdhB* | 1.11E-06 |
| *ΔatpD* | 2.27E-07 |  | *ΔnuoC* | 1.13E-06 |
| *ΔsdhB* | 2.56E-07 |  | *ΔnuoK* | 1.20E-06 |
| *Δpta* | 2.68E-07 |  | *ΔatpE* | 1.21E-06 |
| *ΔatpA* | 3.33E-07 |  | *Δlpd* | 1.71E-06 |
| *ΔnuoK* | 3.33E-07 |  | *ΔfrdC* | 1.73E-06 |
| *ΔsdhD* | 4.29E-07 |  | *ΔsucD* | 1.88E-06 |
| *ΔsucC* | 4.74E-07 |  | *ΔgltA* | 2.56E-06 |
| *ΔgltA* | 6.43E-07 |  | *ΔsucB* | 2.90E-06 |
| *ΔacnB* | 6.47E-07 |  | *Δpta* | 2.97E-06 |
| *ΔfumA* | 7.67E-07 |  | *ΔatpD* | 3.02E-06 |
| *ΔatpC* | 8.00E-07 |  | *ΔatpH* | 3.71E-06 |
| *ΔatpE* | 1.00E-06 |  | *ΔsucA* | 5.32E-06 |
| *ΔadhE* | 1.06E-06 |  | *ΔnuoN* | 5.81E-06 |
| *ΔpykA* | 1.07E-06 |  | *ΔldhA* | 8.71E-06 |
| *ΔglpC* | 1.10E-06 |  | *ΔatpB* | 1.04E-05 |
| *ΔrpiA* | 1.20E-06 |  | *ΔackA* | 1.13E-05 |
| *ΔfumC* | 1.24E-06 |  | *ΔsucC* | 1.67E-05 |
| *Δlpd* | 1.38E-06 |  | *ΔatpA* | 1.96E-05 |
| *ΔfrdC* | 1.38E-06 |  | *ΔadhE* | 2.26E-05 |
| *ΔatpH* | 1.46E-06 |  | *ΔpykA* | 2.29E-05 |
| *ΔfumD* | 1.63E-06 |  | *ΔsdhD* | 4.23E-05 |
| **BW25113 WT** | 1.74E-06 |  | *ΔnapB* | 4.40E-05 |
| *ΔnuoN* | 1.95E-06 |  | *ΔmaeA* | 4.55E-05 |
| *ΔmaeA* | 1.96E-06 |  | *ΔfumB* | 4.64E-05 |
| *ΔnuoF* | 2.04E-06 |  | *ΔphoA* | 5.16E-05 |
| *ΔappC* | 2.28E-06 |  | *ΔnuoB* | 5.50E-05 |
| *ΔatpB* | 2.38E-06 |  | *ΔlacY* | 5.60E-05 |
| *ΔfdoI* | 2.43E-06 |  | *ΔpfkB* | 5.81E-05 |
| *ΔfbaB* | 2.86E-06 |  | **BW25113 WT** | 5.93E-05 |
| *ΔtpiA* | 3.13E-06 |  | *ΔcydB* | 6.11E-05 |
| *Δdld* | 3.30E-06 |  | *ΔglpC* | 6.25E-05 |
| *ΔpflB* | 3.44E-06 |  | *ΔnuoG* | 7.24E-05 |
| *ΔtktA* | 3.67E-06 |  | *ΔacnB* | 7.33E-05 |
| *ΔfdnI* | 3.85E-06 |  | *ΔtdcE* | 7.84E-05 |
| *ΔfumB* | 4.25E-06 |  | *ΔrpiA* | 7.88E-05 |
| *ΔphoA* | 4.44E-06 |  | *Δdld* | 8.33E-05 |
| *ΔpykF* | 5.42E-06 |  | *ΔpflB* | 8.53E-05 |
| *ΔtdcE* | 5.56E-06 |  | *ΔhybB* | 8.86E-05 |
| *Δgnd* | 5.83E-06 |  | *ΔnarY* | 9.23E-05 |
| *ΔpfkB* | 5.86E-06 |  | *ΔglpB* | 9.43E-05 |
| *ΔnuoJ* | 6.00E-06 |  | *ΔlacI* | 9.67E-05 |
| *ΔnuoA* | 6.21E-06 |  | *ΔfumD* | 1.09E-04 |
| *ΔadhP* | 6.25E-06 |  | *ΔhyaB* | 1.13E-04 |
| *ΔnuoG* | 6.45E-06 |  | *ΔadhP* | 1.17E-04 |
| *ΔnuoH* | 6.54E-06 |  | *ΔtktA* | 1.19E-04 |
| *ΔnarZ* | 6.67E-06 |  | *ΔfdnI* | 1.23E-04 |
| *ΔfdnG* | 6.77E-06 |  | *ΔfdoI* | 1.60E-04 |
| *ΔpfkA* | 7.31E-06 |  | *ΔfrdB* | 1.71E-04 |
| *ΔnapB* | 7.60E-06 |  | *ΔnarG* | 2.11E-04 |
| *ΔnarG* | 7.69E-06 |  | *Δeda* | 2.33E-04 |
| *ΔglpA* | 7.73E-06 |  | *ΔtalB* | 2.54E-04 |
| *ΔnrfC* | 8.39E-06 |  | *ΔnapG* | 2.86E-04 |
| *ΔaceB* | 8.52E-06 |  | *ΔppsA* | 3.33E-04 |
| *ΔcydX* | 8.86E-06 |  | *ΔrpiB* | 3.57E-04 |
| *ΔhyaB* | 9.13E-06 |  | *ΔnarZ* | 3.79E-04 |
| *ΔfrdD* | 9.39E-06 |  | *ΔfdnH* | 3.79E-04 |
| *ΔackA* | 9.60E-06 |  | *ΔeutE* | 3.93E-04 |
| *ΔtorC* | 9.72E-06 |  | *ΔhyaA* | 4.29E-04 |
| *ΔglpD* | 1.00E-05 |  | *Δgnd* | 4.33E-04 |
| *ΔdmsC* | 1.00E-05 |  | *ΔnapH* | 4.67E-04 |
| *ΔhyaC* | 1.00E-05 |  | *ΔhyaC* | 4.81E-04 |
| *ΔhycC* | 1.00E-05 |  | *ΔacnA* | 5.00E-04 |
| *ΔdmsB* | 1.04E-05 |  | *ΔaceB* | 5.12E-04 |
| *ΔpoxB* | 1.05E-05 |  | *ΔhycB* | 5.36E-04 |
| *ΔdmsA* | 1.23E-05 |  | *ΔatpG* | 6.07E-04 |
| *ΔfdnH* | 1.25E-05 |  | *ΔfdoH* | 6.15E-04 |
| *ΔlacI* | 1.29E-05 |  | *ΔfbaB* | 6.33E-04 |
| *ΔhybO* | 1.31E-05 |  | *Δccp* | 6.43E-04 |
| *ΔhycE* | 1.33E-05 |  | *ΔtpiA* | 6.52E-04 |
| *ΔnarH* | 1.34E-05 |  | *Δzwf* | 6.67E-04 |
| *ΔeutE* | 1.35E-05 |  | *Δedd* | 7.14E-04 |
| *ΔglpB* | 1.45E-05 |  | *Δicd* | 7.20E-04 |
| *ΔfdoG* | 1.48E-05 |  | *ΔfdnG* | 7.33E-04 |
| *ΔyieF* | 1.48E-05 |  | *ΔglpD* | 7.67E-04 |
| *ΔnapA* | 1.50E-05 |  | *ΔtalA* | 7.74E-04 |
| *ΔwrbA* | 1.52E-05 |  | *ΔyieF* | 7.81E-04 |
| *ΔnapG* | 1.56E-05 |  | *ΔdmsA* | 7.86E-04 |
| *ΔlacY* | 1.61E-05 |  | *ΔnrfC* | 7.93E-04 |
| *ΔfdoH* | 1.62E-05 |  | *ΔhybO* | 8.00E-04 |
| *ΔacnA* | 1.71E-05 |  | *ΔnuoH* | 8.24E-04 |
| *ΔpurT* | 1.84E-05 |  | *ΔhybA* | 8.46E-04 |
| *ΔeutD* | 1.88E-05 |  | *ΔfdoG* | 8.48E-04 |
| *Δzwf* | 1.92E-05 |  | *ΔappB* | 8.61E-04 |
| *ΔhybB* | 1.96E-05 |  | *ΔfrdD* | 8.67E-04 |
| *ΔtorA* | 2.10E-05 |  | *ΔfumE* | 9.12E-04 |
| *ΔnapH* | 2.13E-05 |  | *Δmdh* | 9.13E-04 |
| *ΔhycB* | 2.38E-05 |  | *ΔhycE* | 9.35E-04 |
| *ΔhybA* | 3.08E-05 |  | *Δppc* | 9.41E-04 |
| *ΔfrdB* | 4.05E-05 |  | *ΔpoxB* | 9.62E-04 |
| *ΔcyoA* | 4.07E-05 |  | *ΔnarH* | 1.05E-03 |
| *ΔcydB* | 4.10E-05 |  | *ΔhycC* | 1.06E-03 |
| *ΔtalA* | 4.29E-05 |  | *ΔhycG* | 1.08E-03 |
| *ΔnarY* | 4.55E-05 |  | *ΔdmsC* | 1.08E-03 |
| *ΔhyaA* | 4.62E-05 |  | *ΔnuoE* | 1.08E-03 |
| *ΔfumE* | 4.64E-05 |  | *ΔdmsB* | 1.12E-03 |
| *Δndh* | 5.00E-05 |  | *ΔcydX* | 1.14E-03 |
| *Δppc* | 5.22E-05 |  | *ΔnapA* | 1.27E-03 |
| *Δccp* | 5.31E-05 |  | *ΔhycD* | 1.29E-03 |
| *ΔtktB* | 5.36E-05 |  | *ΔpurT* | 1.32E-03 |
| *ΔppsA* | 5.42E-05 |  | *ΔpykF* | 1.39E-03 |
| *ΔnrfD* | 5.48E-05 |  | *ΔtorC* | 1.42E-03 |
| *ΔhycD* | 5.77E-05 |  | *ΔkefF* | 1.47E-03 |
| *ΔybhA* | 6.00E-05 |  | *ΔglpA* | 1.52E-03 |
| *Δedd* | 6.36E-05 |  | *ΔeutD* | 1.59E-03 |
| *ΔatpI* | 7.14E-05 |  | *ΔkduI* | 1.77E-03 |
| *ΔhycG* | 7.22E-05 |  | *ΔatpI* | 1.92E-03 |
| *ΔkduI* | 7.39E-05 |  | *ΔnuoJ* | 1.93E-03 |
| *ΔappB* | 8.13E-05 |  | *Δndh* | 1.95E-03 |
| *ΔatpG* | 8.33E-05 |  | *ΔnrfD* | 2.16E-03 |
| *ΔnrfA* | 8.64E-05 |  | *ΔyggF* | 2.41E-03 |
| *ΔkefF* | 8.82E-05 |  | *ΔfrdA* | 2.48E-03 |
| *ΔcyoB* | 1.08E-04 |  | *ΔtktB* | 2.56E-03 |
| *Δpck* | 1.14E-04 |  | *ΔwrbA* | 2.60E-03 |
| *Δmdh* | 1.23E-04 |  | *ΔhybC* | 2.74E-03 |
| *Δicd* | 1.24E-04 |  | *ΔnuoL* | 3.53E-03 |
| *ΔyggF* | 1.25E-04 |  | *Δpck* | 3.59E-03 |
| *ΔtalB* | 1.35E-04 |  | *ΔnrfA* | 3.70E-03 |
| *ΔfrdA* | 1.39E-04 |  | *ΔaceF* | 3.86E-03 |
| *ΔaceA* | 1.44E-04 |  | *ΔnuoF* | 4.23E-03 |
| *ΔcyoD* | 1.47E-04 |  | *ΔnuoA* | 4.83E-03 |
| *ΔhybC* | 1.52E-04 |  | *ΔputA* | 5.33E-03 |
| *Δpgi* | 1.60E-04 |  | *ΔtorA* | 5.65E-03 |
| *ΔputA* | 1.63E-04 |  | *ΔglcB* | 6.00E-03 |
| *Δfbp* | 1.64E-04 |  | *ΔybhA* | 6.52E-03 |
| *ΔcyoC* | 1.67E-04 |  | *ΔaceA* | 7.83E-03 |
| *ΔnarI* | 1.79E-04 |  | *ΔglpX* | 1.08E-02 |
| *ΔnuoE* | 1.90E-04 |  | *Δfbp* | 1.20E-02 |
| *Δeda* | 1.95E-04 |  | *Δpgi* | 1.25E-02 |
| *ΔglcB* | 2.00E-04 |  | *ΔnarI* | 1.25E-02 |
| *ΔglpX* | 4.33E-04 |  | *Δrpe* | 1.44E-02 |
| *ΔnuoL* | 5.22E-04 |  | *ΔcyoD* | 2.31E-02 |
| *ΔrpiB* | 8.24E-04 |  | *ΔcyoA* | 2.80E-02 |
| *ΔaceF* | 1.25E-03 |  | *ΔcyoC* | 3.00E-02 |
| *Δrpe* | 1.50E-03 |  | *ΔcyoB* | 3.04E-02 |
