## Supplementary File 5 for "UNRAVELING CRP/cAMP-MEDIATED METABOLIC REGULATION IN *ESCHERICHIA COLI* PERSISTER CELLS"

**Supplemental File 5. The knockout strains generated using the *E. coli* K-12 MG1655 background in this study.** Their functions and association with CRP/cAMP regulation and ampicillin or ofloxacin persister formation in *E. coli* organisms reported in the literature were described (not a comprehensive review). Empty boxes indicate a lack of information. We observed that the proteins associated with the genes below were predominantly downregulated in the Δ*crp* strain in our study, in line with the literature.

| **Gene name** | **Function** | **Under CRP/cAMP regulation** | **Persistence in *E. coli*** |
| --- | --- | --- | --- |
| Δ*crp* | DNA-binding transcriptional dual regulator CRP | Positively and negatively regulated (Gonzalez-Gil, 1998; Zheng, 2004) | Decrease in the late stationary phase (Mok et al., 2015) |
| Δ*cyaA* | Adenylate cyclase | Negatively regulated (Aiba, 1985) | Increase in the exponential phase (Chu et al., 2012; Molina-Quiroz et al., 2018; Sulaiman & Lam, 2020; Yamasaki et al., 2020) |
| Δ*sucA*  Δ*sucB*  Δ*sucC* | 2-oxoglutarate decarboxylase, thiamine-requiring | Positively regulated (Gosset et al., 2004; Khankal et al., 2009; Lynch & Lin, 1996; Salgado et al., 2013; Surmann et al., 2020; Tsai et al., 2018; Zhang et al., 2005; Zheng, 2004) | Decrease in the lag phase or the stationary phase (Liu et al., 2017; Ma et al., 2010; Orman & Brynildsen, 2015)  Increase in exponential phase at low ampicillin concentrations (Kohanski et al., 2007)  Decrease in the lag phase (Luidalepp et al., 2011) |
| Δ*lpd* | Lipoamide dehydrogenase | Positively and negatively regulated (Cunningham & Guest, 1998; Zhang et al., 2005) |  |
| Δ*sdhA* | Succinate:quinone oxidoreductase, FAD binding protein | Positively regulated (Lynch & Lin, 1996; Surmann et al., 2020; Tsai et al., 2018; Zhang et al., 2005; Zheng, 2004) | Decrease in the exponential phase (Mohiuddin et al., 2022)  Decrease in the lag phase (Leatham-Jensen et al., 2016) |
| Δ*gltA* | Citrate synthase | Positively regulated (Gosset et al., 2004; Lynch & Lin, 1996; Perrenoud & Sauer, 2005; Salgado et al., 2013; Surmann et al., 2020; Tsai et al., 2018; Zhang et al., 2005; Zheng, 2004) |  |
| Δ*aceE* | Pyruvate dehydrogenase | Positively regulated (Quail et al., 1994; Zhang et al., 2005) |  |
| Δ*mdh* | Malate dehydrogenase | Positively regulated (Khankal et al., 2009; Perrenoud & Sauer, 2005; Salgado et al., 2013; Vogel et al., 1987) | Decrease in the lag phase (Orman & Brynildsen, 2015)  Increase or no change depending on the growth phase (Luidalepp et al., 2011) |
| Δ*nuoI* | NADH:quinone oxidoreductase subunit I |  |  |
| Δ*nuoM* | NADH:quinone oxidoreductase subunit M |  |  |
| Δ*atpA* | ATP synthase F_1_ complex subunit α |  | Decrease in the exponential phase (Mohiuddin et al., 2022)  Decrease in the exponential phase (Lobritz et al., 2015) |
| Δ*atpB* | ATP synthase F_o_ complex subunit a | Positively regulated (Salgado et al., 2013) |  |
| Δ*atpC* | ATP synthase F_1_ complex subunit ε |  | Decrease in the lag phase (Orman & Brynildsen, 2015) |
| Δ*atpD* | ATP synthase F_1_ complex subunit β | Positively regulated (Salgado et al., 2013) | Decrease in the exponential phase (Mohiuddin et al., 2022) |
| Δ*frdC* | Fumarate reductase membrane protein FrdC |  |  |
| Δ*pgi* | Glucose-6-phosphate isomerase |  |  |
| Δ*zwf* | NADP^+^-dependent glucose-6-phosphate dehydrogenase |  | No effect (Leatham-Jensen et al., 2016) |
| Δ*tktB* | Transketolase 2 |  |  |
| Δ*talA* | Transaldolase A |  |  |
