## Supplementary File 6 for "UNRAVELING CRP/cAMP-MEDIATED METABOLIC REGULATION IN *ESCHERICHIA COLI* PERSISTER CELLS"

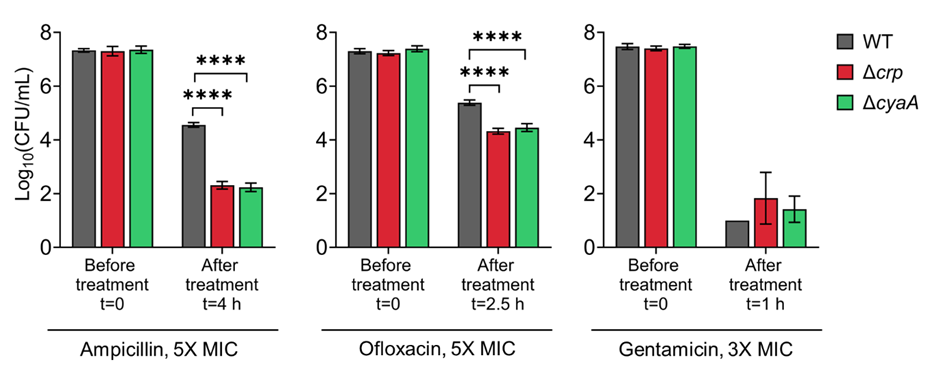

**Supplemental File 6. Persister levels of *E. coli* K-12 MG1655 WT, Δ*crp*, and Δ*cyaA* strains in late stationary phase.** Cells were treated with ampicillin (5× MIC for 4 h), ofloxacin (5× MIC for 2.5 h), and gentamicin (3× MIC for 1 h). Concentrations and treatment durations were selected based on (Zeng et al., 2022). n=4. Statistical significance was observed between control and mutant strains (****P < 0.0001, One-way ANOVA with Dunnett’s multiple comparisons test). The data for each time point represent the mean value ± standard deviation.
