## Supplementary File 7 for "UNRAVELING CRP/cAMP-MEDIATED METABOLIC REGULATION IN *ESCHERICHIA COLI* PERSISTER CELLS"

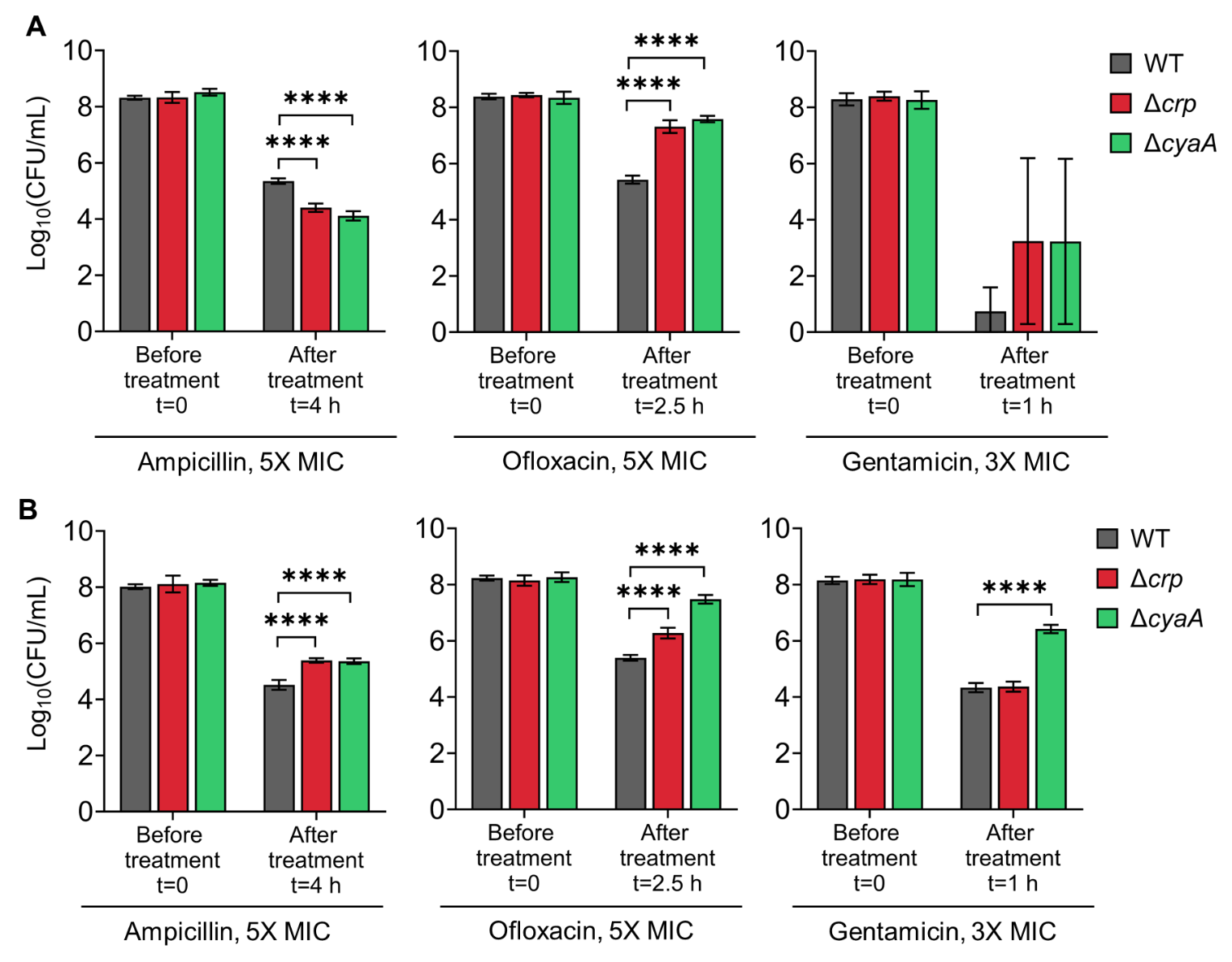

**Supplemental File 7. Persister levels of *E. coli* K-12 MG1655 (A) and BW25113 (B) WT, Δ*crp*, and Δ*cyaA* strains in the exponential growth phase.** Cells were treated at mid-exponential phase (OD₆₀₀ ~0.25) with ampicillin (5× MIC for 4 h), ofloxacin (5× MIC for 2.5 h), and gentamicin (3× MIC for 1 h). Treatment concentrations and durations were based on conditions described in (Zeng et al., 2022). n=4. Statistical significance was observed between control and mutant strains (****P < 0.0001, One-way ANOVA with Dunnett’s multiple comparisons test). The data for each time point represent the mean value ± standard deviation.
